# SAMURAI: Shallow Analysis of copy nuMber alterations Using a Reproducible And Integrated bioinformatics pipeline

**DOI:** 10.1101/2024.09.30.615766

**Authors:** Sara Potente, Diego Boscarino, Dino Paladin, Sergio Marchini, Luca Beltrame, Chiara Romualdi

**Author notes:** Co-last authors (Luca Beltrame, Chiara Romualdi). **Correspondence to:** Luca Beltrame, IRCCS Humanitas Research Hospital.

## Abstract

Shallow whole-genome sequencing (sWGS) offers a cost-effective approach to detect copy number alterations (CNAs). However, there remains a gap for a standardized workflow specifically designed for sWGS analysis. To address this need, in this work we present SAMURAI a bioinformatics pipeline specifically designed for analyzing CNAs from sWGS data in a standardized and reproducible manner.

SAMURAI is built using established community standards, ensuring portability, scalability, and reproducibility. The pipeline features a modular design with independent blocks for data pre-processing, copy number analysis, and customized reporting. Users can select workflows tailored for either solid or liquid biopsy analysis (e.g., circulating tumor DNA), with specific tools integrated for each sample type. The final report generated by SAMURAI provides detailed results to facilitate data interpretation and potential downstream analyses.To demonstrate its robustness, SAMURAI was validated using simulated and real-world data sets. The pipeline achieved high concordance with ground truth data and maintained consistent performance across various scenarios.

By promoting standardization and offering a versatile workflow, SAMURAI empowers researchers in diverse environments to reliably analyze CNAs from sWGS data. This, in turn, holds promise for advancements in precision medicine.

## Introduction

DNA copy number alterations (CNAs) are defined as gains or losses of DNA segments (at least 50 bp long) [1]. They are distinct from copy number variations (CNVs) that are smaller in length and often inherited [2].

CNAs and somatically acquired CNAs (SCNAs) can contribute to different types of diseases, such as schizophrenia, Crohn’s disease, developmental diseases and many others [3]. In addition, CNAs have very important roles in cancer, which are frequently characterized by genomic instability [4–6].

Recently, more insights on CNAs and SCNAs have been made possible through the use of next generation sequencing (NGS) technologies. In particular, shallow whole genome sequencing (sWGS), that is sequencing of the whole genome at low depth (from 1X to as low as 0.1X), provided to be a viable and cost-effective solution towards the investigation of CNAs in cancer and other pathologies [4,7–9]. Additionally, sWGS also proved effective in the identification of CNAs in biological fluids [5,10,11], such as routine noninvasive prenatal testing (NIPT) [12] or as a potential approach to monitor cancer patients or evaluate therapeutic options [5,13].

Over the course of the years, the bioinformatics community has developed several approaches for the analysis of CNAs with sWGS data, such as QDNAseq [7], WisecondorX [9], ichorCNA [10,14] and others [15,16] on both formalin-fixed paraffin-embedded (FFPE) tumor tissues or bodily fluids. These are stand-alone tools, which are often integrated into custom bioinformatics pipelines, which are tailored for specific use-cases. This is a significant challenge for standardization and reproducibility of the results produced with these pipelines.

To tackle this problem, standards such as the workflow description language (WDL) [17] or the common workflow language (CWL) [18] along with workflow managers such as Snakemake [19] and Nextflow [20] have been developed. In particular, the Nextflow-derived nf-core [21,22] offers a vast collection of bioinformatics pipelines, but at the time of writing no specific workflow suited for the analysis of sWGS data exists. To address this need, we have developed the first nf-core based pipeline to process sWGS data, SAMURAI (Shallow Analysis of copy nuMber alterations Using a Reproducible And Integrated bioinformatics pipeline). SAMURAI integrates different methods for pre-processing data, performing CNA analysis, along with optional post-processing steps, leveraging the nf-core standards and vast array of pre-made analysis modules. SAMURAI is freely downloadable at https://github.com/DIncalciLab/samurai and distributed under a free and open source (FOSS) license (MIT).

## Materials and Methods

### Implementation

SAMURAI has been implemented by using Nextflow workflow management system [20] and adhering to the existing nf-core guidelines for developers [21,22].

### Pipeline Architecture

SAMURAI is a reproducible pipeline for detecting copy number variations in low-pass or ultra low-pass whole genome sequencing data. The pipeline incorporates several tools from the bioinformatics community, containerized (either Docker [23] or Singularity / Apptainer [24]) to ensure the maximum reproducibility. The workflow of the pipeline can be divided into three main areas: pre-processing, copy number detection, and post-processing (Figure 1).

**Figure 1.**
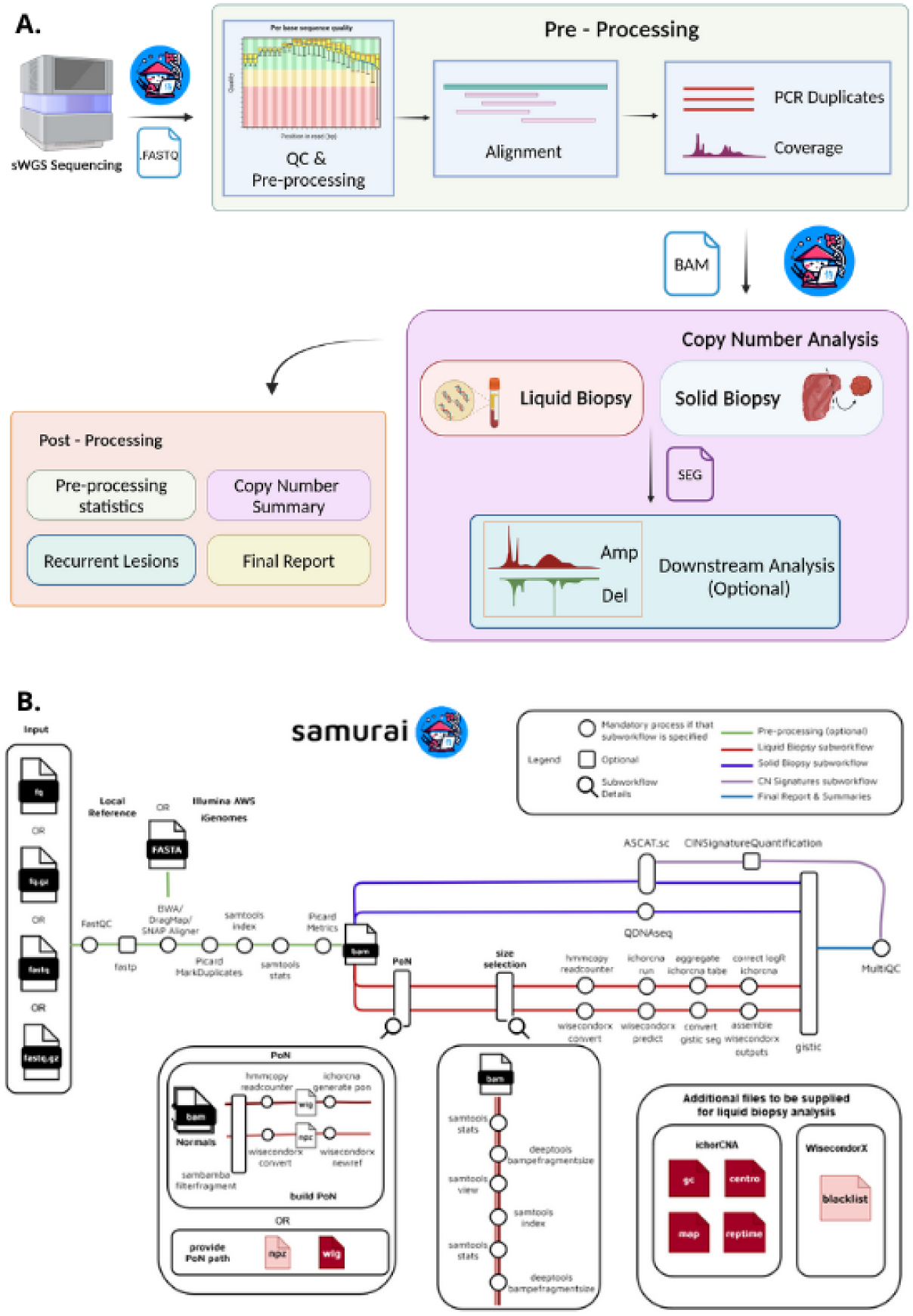
Overview of the SAMURAI pipeline. **A)** General workflow illustrating the three main stages: pre-processing, copy number detection, and post-processing. **B)** Detailed breakdown of the pipeline’s architecture, representing modules and subworkflows responsible for each stage of the analysis.

### Pre - processing

The pre-processing phase of SAMURAI includes a first step of quality control on the input data, followed by alignment to a reference genome. The entire pre-processing phase can be skipped when pre-aligned data is supplied.

The list of software included in SAMURAI’s pre-processing phase is reported in *Table 1*.

**Table 1.** Detailed list of software included in SAMURAI pre-processing block.

| Tool | Input | Subworkflow | Ref. |
| --- | --- | --- | --- |
| FastQC | FASTQ | QC, quality trimming adapter removal | [25] |
| fastp | FASTQ |  | [26,27] |
| BWA INDEX | FASTA | Reference genome indexing | [28,29] |
| BWA-MEM | FASTQ, FASTA | Alignment & marking PCR duplicates | [28] |
| BWA-MEM2 |  |  | [29] |
| DragMap |  |  | [30] |
| SNAP aligner |  |  | [31] |
| Samtools | BAM |  | [32] |
| Picard | BAM |  | [33] |

#### Quality trimming and adapter sequence removal (optional)

Starting with raw sequence data, SAMURAI performs quality control checks with FastQC [25]. If required, adapter sequences removal and quality trimming can be performed with fastp [26]. Additionally, SAMURAI can process unique molecular identifiers (UMIs) if the library design includes them.

#### Alignment to reference genome

In case raw sequence data are supplied, SAMURAI will perform alignment against a reference genome with two different algorithms (BWA-MEM and BWA-MEM2). Reference genomes can be user-supplied, taken from Illumina’s iGenomes, or using the refgenie reference manager [34]. The pipeline will also create an aligner index in case it is missing. The alignment step is also coupled with removal of PCR duplicates.

### Copy Number Analysis

The core of the SAMURAI pipeline is copy number calling. It offers two types of analysis workflow depending on the nature of the samples:one catering to the processing of tissues (“solid biopsy”) and another for the identification of aberrant CNAs in other biological fluids (“liquid biopsy”). This design allows users to specify the workflow based on their specific needs. The main tools used by both workflows are indicated in Table 2.

**Table 2.** Summary of tools included in SAMURAI copy number analysis block.

| **Solid Biopsy workflow** |  |  |  |
| --- | --- | --- | --- |
| **Tool** | **Input** | **Description** | **Ref.** |
| QDNAseq | BAM | Generates copy number segments, correcting for sequence bias (GC content, mappability) | [7] |
| ASCAT.sc | BAM | Calls absolute copy numbers. It also estimates tumor purity and sample ploidy. | [35] |
| CINSignatureQuantification | SEG | Quantifies chromosomal instability signatures based on absolute copy number values. | [8] |
| **Liquid Biopsy workflow** |  |  |  |
| **Tool** | **Input** | **Description** | **Ref.** |
| Samtools | BAM | Enhances ctDNA copy number calling performance by performing size selection on ctDNA. | [32,36] |
| ichorCNA | BAM,<br>Wiggle | Calls absolute copy number values in ctDNA samples and estimates tumor fraction. | [10] |
| WisecondorX | .npz | Identifies regions with altered copy number compared to a panel of normals and calculates a copy number profile abnormality (CPA) score for each sample. | [9,11] |

The different steps are interconnected through the use of custom scripts provided with the pipeline to harmonize and integrate analysis outputs.

#### Solid Biopsy workflow

The solid biopsy workflow (*Figure 2*) processes data by binning, correcting for sequence biases such as GC content and mappability, and then segments of identical copy number. Depending on the needs of the user, two different tools can be used, either to generate log2 ratios of observed vs expected reads (QDNAseq) [7] or to call absolute copy numbers (ASCAT.sc) [35].

**Figure 2.**
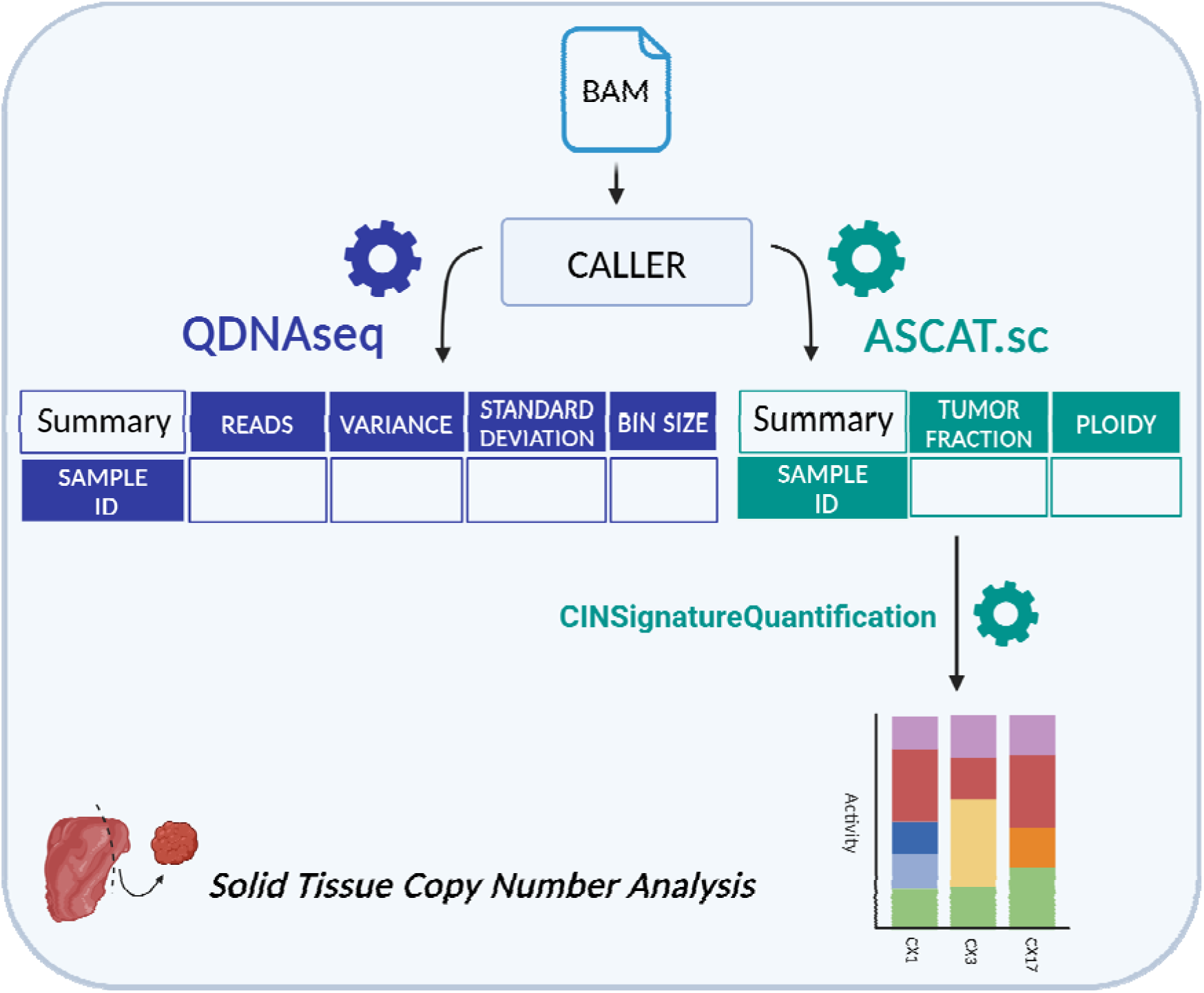
Schema of solid biopsy analysis workflow, including the types of data collected for the final report.

QDNAseq does not produce absolute CNA calls itself, but its results can be used as input for downstream analyses, both within SAMURAI and with external tools like shallowHRD [37], ACE [38] or RASCAL [15]. ASCAT.sc, an evolution of the ASCAT algorithm originally developed for microarrays, is used to estimate absolute copy number values from the supplied aligned data, and, in case of tumor samples, also estimate their purity and ploidy.

Both tools generate plots showing the state of genome-wide copy number profiles for each analyzed sample, which are then included in SAMURAI’s final output files (an example report is available as Supplementary File 1).

#### Liquid Biopsy workflow

In addition to the analysis of solid samples, SAMURAI features a complementary workflow specifically designed for analyzing CNAs in cell-free DNA isolated from bodily fluids.

This workflow, summarized in *Figure 3*, uses specialized tools designed for this purpose: ichorCNA [14] and WisecondorX [9].

**Figure 3.**
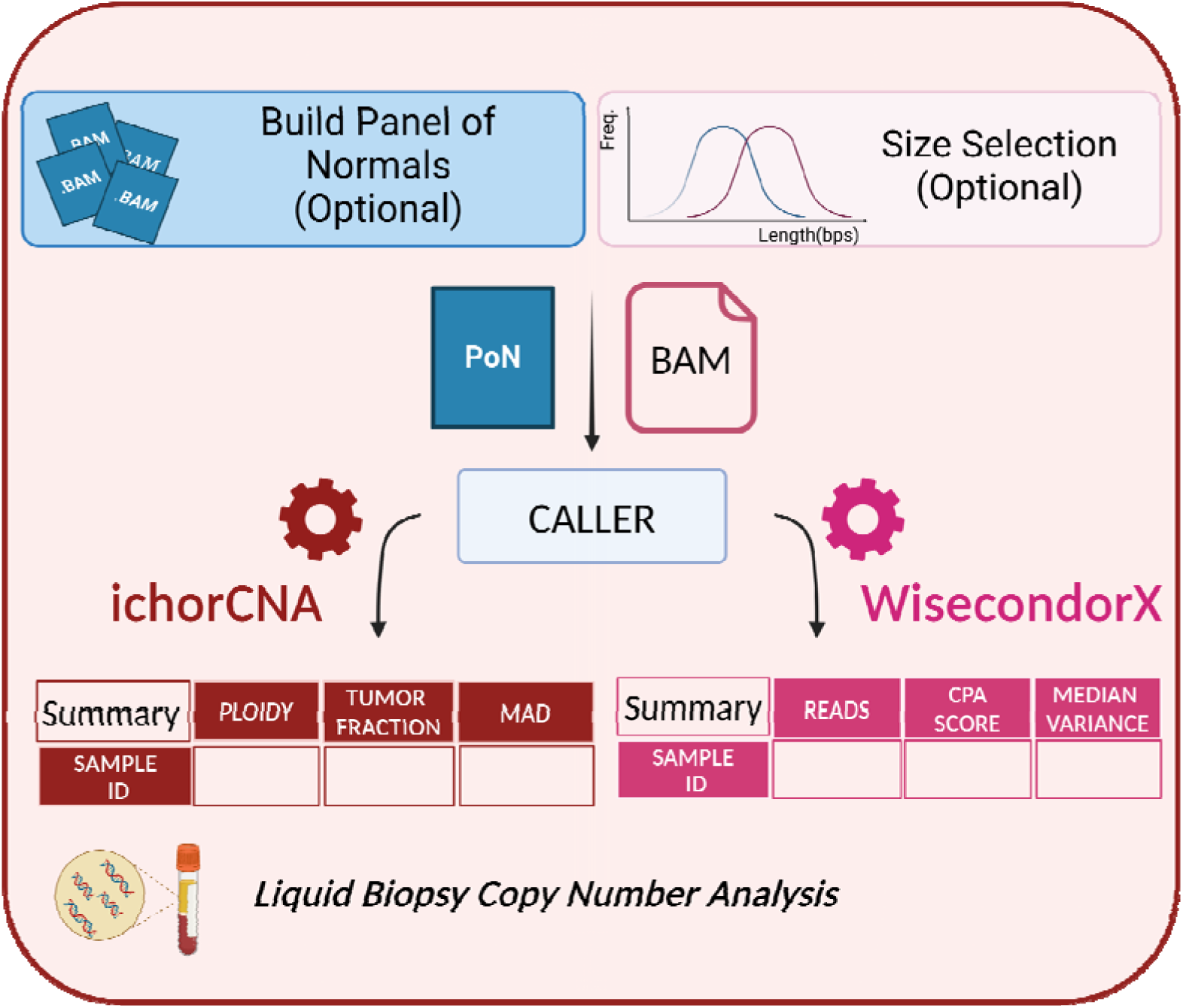
Schema of liquid biopsy analysis workflow, with the output tables that are included in the final report.

Leveraging the knowledge that ctDNA fragments are typically shorter than healthy cfDNA [36], SAMURAI offers an optional size selection subworkflow. This subworkflow preferentially enriches for ctDNA fragments by selecting those within a specific size range, typically between 90 and 150 base pairs, potentially improving the accuracy of CNA detection in ctDNA analysis.

Similar to the solid biopsy analysis, depending on the needs of the user, two tools can be used to call absolute copy numbers (ichorCNA [10]) or to identify regions with altered copy numbers (WisecondorX [9]).

Both tools rely on a panel of normals (PoN) for normalization and to improve accuracy of CNA calls. SAMURAI provides users with the flexibility to either construct the PoN internally during the analysis, or incorporate a previously-built panel.

ichorCNA [10] estimates absolute copy number and, in case of tumor samples, estimates purity (expressed as tumor fraction, TF) and ploidy. WisecondorX [9] identifies regions of gains and losses and computes a copy number abnormality score (CPA) [11], an indication of the overall genomic instability of the sample.

In a similar fashion to the solid biopsy workflow, both tools included in the liquid biopsy analysis generate plots depicting the state of these copy number profiles for each analyzed sample. These plots are then included in SAMURAI’s final output (an example of the liquid biopsy analysis is provided as Supplementary File 2).

#### Downstream Analysis of Copy Number Alterations

##### Chromosomal instability (CIN) signature extraction

To understand chromosomal instability (CIN) patterns, SAMURAI leverages a framework by Drews et al. [39] that analyzes absolute copy number profiles and identifies 17 distinct pan-cancer CIN signatures. By incorporating this framework, SAMURAI presents a matrix of normalized signature activities alongside a bar plot summarizing these activities, allowing users to easily interpret the CIN landscape within the report.

##### Identification of recurrent copy number alterations

SAMURAI also aids in identifying frequently recurring copy number changes (SCNAs) in cancer samples. It integrates GISTIC 2.0 [40] software to pinpoint these recurrent regions across multiple samples and localize any target genes within them. SAMURAI then transforms GISTIC’s output into a user-friendly format, providing tables and plots that clearly represent these identified SCNAs and associated genes.

### Post-Processing

To aggregate results from the different tools comprehensively, SAMURAI incorporates outputs from each phase into a final report generated with MultiQC [41]. This report includes information on quality control, providing an overview about sequencing run and pre-processing, aiding also in identifying potential problematic samples. SAMURAI also provides users with custom tables summarizing the copy number analysis section, which may include tumor fraction, ploidy values for each sample, or measures of copy number instability such as CPA from WisecondorX analysis. Additionally, SAMURAI also includes specific plots, such as recurrent alterations plots or signature activities plots, to assist in the interpretation of the results.

Alongside the final report, the pipeline stores outputs and versions of different software tools in the output folder chosen by the user.

### Analysis data sets

To develop and test the functionality of SAMURAI, we used three different data sets:

- A diluted data set from a public, previously published artificial sample [16] (hereafter called data set S)
- A set of 218 sWGS samples from 205 Stage I ovarian cancer [4], EGA ID EGAS00001004961 (hereafter called data set T)
- A set of 12 plasma samples withdrawn at time of diagnosis from 12 ovarian cancer patients part of a larger and previously published cohort [5], EGA ID EGAS00001004670 (hereafter called data set P)

### Simulated data downsampling

To simulate real-world coverage scenarios, data set S was downsampled with Picard DownSampleSam [33] to simulate different coverages ( 0.1X, 0.3X, 0.5X, and 0.7X).

For those analysis methods which relied on a panel of normals, we created 30 simulated control samples by downsampling normal sample SM-74-NEG from GATK test data (https://42basepairs.com/browse/s3/gatk-test-data/cnv/somatic) at identical coverages (*Supplementary Table S2)* using SAMtools [32], Sambamba [42] and Bedtools [43]. Script used for download data and downsample is available at https://github.com/DIncalciLab/SAMURAI_paper_scripts.

Final average and median coverages of simulated samples were computed with mosdepth [44], including bases with 0 coverage.

### Benchmarking of copy number calls with simulated data

To compare results from SAMURAI to the ground truth provided in the original publication [16] for their simulated sample, the genomic coordinates from the ground truth were lifted over from hg19 to the hg38 assembly using https://genebe.net/tools/liftover. The R package CNVMetrics [45,46] was used to compute the overlap (Szymkiewicz-Simpson coefficient) similarity score between inferred and ground truth copy numbers. We made the comparisons for results from ASCAT.sc, ichorCNA and WisecondorX.

### Benchmarking of real-world data sets

Data set T was analyzed with SAMURAI starting from pre-aligned files (BAM; hg38 genome), using the solid biopsy pipeline. ASCAT.sc was run to obtain absolute copy number values, using a bin size of 30 Kbp. After copy number calling, overlap (Szymkiewicz-Simpson coefficient) similarity score was computed to evaluate the concordance between the inferred and reference copy number segments. Additionally, proportions of gains and losses were computed on 30 kbp bins on chromosome 8 for high grade serous (HGSOC) histotype with the function cnFreq() of the GenVizR package [47] and the Pearson correlation coefficient [48] was computed between the two proportions with the R function [49] *cor()*.

Analysis of data set P was carried out with SAMURAI using the liquid biopsy pipeline, ichorCNA as copy number caller and a 500 Kbp bin size. Prior to copy number calling, library target size selection was performed with the appropriate option in SAMURAI to increase sensitivity and specificity [36]. The initial normal contamination states of ichorCNA were set ranging from 0.9 to 0.999 instead of the default to increase sensitivity. A set of 11 healthy controls from the original publication [5] was used as a panel of normals (PoN). Tumor fraction estimates from SAMURAI were compared to those from the original data set. To evaluate the concordance between the two analyses, we computed the Pearson correlation coefficient between SAMURAI and the fractions from the original work. Correlation plots for both analyses were obtained from the ggpubr R package [50].

## Results

### Benchmarking SAMURAI

After developing SAMURAI, our aim was to evaluate the pipeline’s performance both in terms of the entire workflow and from a biological perspective in order to verify reliably and reproducibly. To this aim, we conducted tests using both simulated data (data set S; <u>Materials and Methods</u>) and two real data sets, a previously published sWGS analysis of biopsies withdrawn from patients with early-stage epithelial ovarian cancer (data set T) and a data set with copy number data obtained from plasma samples withdrawn from patients with late-stage high grade serous ovarian cancer (data set P).

#### Case Study 1 : Evaluation of SAMURAI on simulated data

We used a synthetic data set (data set S; see <u>Materials and Methods</u>) to test both solid biopsy and liquid biopsy sub-workflows within SAMURAI, setting different bin sizes to simulate real-world analysis scenarios (50 kbp and 500 kbp, respectively). All the tools used in the copy number analysis were run with default parameters.

We compared the inferred copy-number alterations from ASCAT.sc, ichorCNA, and WisecondorX with the ground truth CNAs for the original A1 sample using the Szymkiewicz-Simpson metric (overlap coefficient; Materials and Methods). All tools achieved a high degree of similarity, with an overlap coefficient greater than 0.9 for gains and 0.8 for losses (*Table 3*), indicating strong agreement between the inferred CNAs obtained using different tools in SAMURAI and the true CNAs.

**Table 3.** Szymkiewicz-Simpson coefficients for gains and losses between original sample A1 and the ground truth sample.

| Caller | Gain | Loss |
| --- | --- | --- |
| ASCAT.sc | 0.99 | 0.99 |
| ichorCNA | 0.91 | 0.98 |
| WisecondorX | 0.99 | 0.82 |

To further evaluate real-world scenarios, we diluted the original samples at different coverages (*Supplementary Table S1*), as well as synthetic normal samples used to build the panel of normals (PoN) for the simulation of liquid biopsy workflows (*Supplementary Table S2*), and we evaluated the consistency of copy number calling across the random downsampling. Similar performance was observed for *in silico* diluted samples (*Supplementary Figure S1*). The overlap coefficient remained high for gains ( > 0.8) and losses (> 0.7) in the diluted samples compared with the ground truth.

#### Case Study 2 : Solid Biopsy Analysis on Ovarian Cancer

In order to fully evaluate SAMURAI’s capability to extract biologically meaningful results, we ran the pipeline over a previously published data set of 218 biopsies withdrawn from 204 patients with early-stage epithelial ovarian cancer (EOC; [4]). This data set contains samples from the five major histotypes of EOC (high grade serous, HGSOC; low grade serous, LGSOC; endometrioid, EC; clear cell, OCCC; mucinous, MOC) and is ideal for evaluating the pipeline as the various histotypes exhibit distinct copy number patterns.

After estimating absolute copy number values, we evaluated the concordance between the previously published result and SAMURAI output by computing the overlap coefficient in a similar fashion as the simulated data (<u>Materials and Methods</u>). The median values for all histotypes were all above 0.8, indicating a large agreement with the previously published results (Table 4). Similar results were observed in the majority of samples when analyzing samples individually. SAMURAI correctly did not call alterations in samples without any gains or losses, such as sample 105 (EC histotype) and sample 156 (LGSOC histotype; *Supplementary Table S3*). Gains and losses included previously reported frequent alterations, such as copy number gains in HGSOC affecting the q arm of chromosome 8 [51,52], as shown in Figure 4. Correlation between previously published data and the results from SAMURAI in this representative case was high (Pearson’s correlation coefficient 0.99; Figure 4 and *Supplementary Figure S2*).

**Figure 4.**
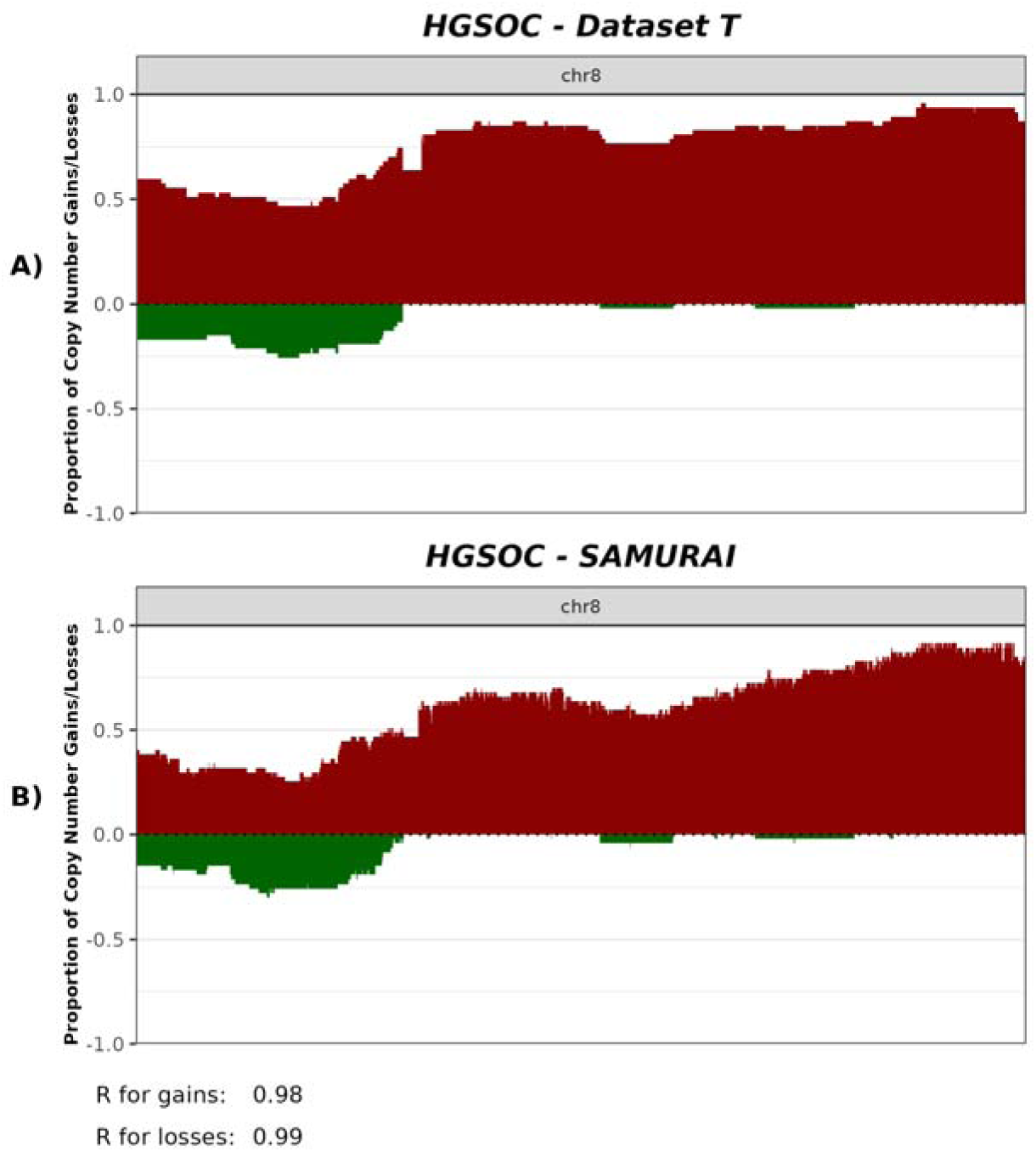
Proportions of gains and losses on chromosome 8 of HGSOC samples in data set T. **A)** Proportions of gains and losses from the results of the original work. **B)** Proportion of gains and losses in SAMURAI. On the Y-axis are represented proportions of alteration; the X-axis represent the genomic coordinates on chromosome 8. R: Pearson’s correlation coefficient.

**Table 4.** Median values of pairwise overlap coefficient for each sample in data set T, divided by histotype. The coefficient was computed for both gains and losses.

| Histotype | Median Coeff. GAINS | Median Coeff. Losses |
| --- | --- | --- |
| HGSOC | 0.94 | 0.95 |
| LGSOC | 0.85 | 0.94 |
| EC | 0.92 | 0.95 |
| MOC | 0.93 | 0.92 |
| OCCC | 0.88 | 0.94 |

#### Case Study 3: Liquid Biopsy Analysis on Ovarian Cancer

In order to test SAMURAI’s liquid biopsy workflow on real-world data, we retrieved sequencing data from 12 plasma samples from 12 distinct late-stage HGSOC patients (withdrawn at time of diagnosis), part of a previously published study (data set P), and then ran SAMURAI with ichorCNA to estimate CNAs and tumor fraction (see Materials and Methods).

The estimates of tumor fraction made by SAMURAI were largely comparable with the previous study (Table 4). Pearson’s correlation analysis comparing the tumor fraction (TF) from the previously published results with the TF calculated by SAMURAI was above 0.9 (0.95; *Supplementary Figure S3*). Analysis of recurrent alterations made by SAMURAI identified two significant regions with copy number gains (3q26.2 and 8q24.21) previously described with HGSOC.

**Table 5.** ichorCNA tumor fraction (TF) estimates from SAMURAI compared to the original values from data set P.

| Sample | TF SAMURAI | TF data set P |
| --- | --- | --- |
| 21531-PL1 | 12.91% | 10.84% |
| 21553-PL1 | 4.19% | 5.75% |
| 21557-PL1 | 8.45% | 7.79% |
| 21564-PL1 | 9.61% | 6.63% |
| 21566-PL1 | 13.58% | 13.66% |
| 21569-PL1 | 18.54% | 18.6% |
| 21572-PL1 | 9.22% | 7.7% |
| 21580-PL1 | 14.51% | 14.76% |
| 21611-PL1 | 13.32% | 12.86% |
| 21614-PL1 | 4.44% | 5.81% |
| 21624-PL1 | 10.86% | 10.97% |
| 21627-PL1 | 6.01% | 6.13% |

#### SAMURAI runtime and memory usage

A typical run of SAMURAI on an HPC platform took approximately 1 hour and 30 minutes for two hundred sWGS samples at 0.5X with the ‘solid biopsy’ workflow (data set T), using approximately 30G of memory for the most compute intensive task (alignment). In the case of the liquid biopsy workflow (data set P), analysis of pre-aligned BAM files (coverage between 1 and 1.5X) took 15 minutes with a peak memory usage of 3G.

Successful analysis steps are cached, so subsequent re-runs only execute those steps which have changed from the previous run.

## Discussion

We have developed SAMURAI, a standardized and reproducible pipeline for the analysis of low pass whole-genome sequencing data. The field of shallow whole-genome sequencing has shown many useful practical applications, like non-invasive prenatal testing (NIPT) [12], cancer diagnostics applied to different sample matrices, such as liquid biopsies [53–55] or formalin-fixed, paraffin-embedded (FFPE) [56] samples with diagnostic and/or prognostic purposes. However, although established approaches exist for whole-genome and whole-exome sequencing [47], most of the analysis methods employed for sWGS rely on “ad hoc” approaches specific for one particular application, or even for a specific research group’s needs. Given the potential benefit of sWGS-based approaches in the clinic [14,57,58], it is imperative that analysis tools are built to be both reliable and reproducible. Thus, SAMURAI aims to fill this gap, by taking advantage of the pre-existing nf-core [21,22] resource to build a robust pipeline. By leveraging the nf-core framework [21,22], SAMURAI inherits its scalability, enabling execution across various computational environments, from local machines to high-performance computing clusters and cloud platforms, and add-ons such as the Seqera Platform exist to simplify execution and configuration for those users which are unfamiliar with the framework. The pipeline prioritizes flexibility by incorporating optional subworkflows tailored to specific needs of the users. Moreover, some of these workflows can be particularly relevant for cancer analysis, such as size selection for ctDNA fragments enrichment, improving the accuracy of CNAs detection, or the identification of recurrent CNAs across a cohort of patients and identifying potential target genes among them. By employing widely used bioinformatics tools for copy number analysis and container technologies for ensuring modular and immutable software in each step, SAMURAI adheres to established bioinformatics standards that maintain a reliable and reproducible analysis environment while offering customizable workflows.

We evaluated SAMURAI’s robustness using simulated data (data set S). The analysis yielded very high copy number concordance (> 80%) compared to the ground truth, indicating strong agreement with actual alterations. We further tested SAMURAI’s performance under various simulated dilution levels to reflect real-world scenarios, showing an overall consistent, robust and reliable performance.

While simulated data provides a controlled environment for testing, it is of course clear that simulations cannot fully represent real-world data. For this reason we also tested SAMURAI with two additional data sets: a large cohort of tumor tissue biopsies [4] and a selection of circulating free DNA (cfDNA) extracted from plasma of ovarian cancer patients [5]. When analyzing data set T, a collection of copy number data from early-stage epithelial ovarian cancer [4], we observed high concordance in the results with the previously published data. The differences in CNA calling compared to the original work can be attributed to two differences: firstly, in the original publication [4], they integrated variant calling with a second sequencing run to estimate the purity / ploidy of samples, while sWGS alone cannot be used to reliably call variants; secondly, they used ACE [38] for purity / ploidy estimation, while SAMURAI uses ASCAT.sc, and the two methods are deeply different in how they calculate their estimations. Despite these differences, it is important to highlight that our analysis showed high concordance between SAMURAI and the ground truth method. A small number of samples were unable to be analyzed properly and reported a purity of 1 and a ploidy of 2: these were either very low purity samples (only analyzable with additional data such as variant calling information), or samples with no detectable alterations [5].

Given the rise in use of liquid biopsy for tumor detection, monitoring, and prognostic purposes [52,59,60], it is essential for a well-constructed pipeline to be able to analyze ctDNA samples correctly and reproducibly. For this reason we used data set P to benchmark the capabilities of SAMURAI in this setting, using ichorCNA. Our results closely matched the original data in terms of detected tumor fraction, with a correlation close to 95%. This was achieved also in samples which were particularly problematic due to low cfDNA concentration. Characteristic alterations of the pathology, for example copy number gains on chromosome 8q24.3 were also detected [4].

SAMURAI offers several advantages over custom-made pipelines. Firstly, it packages current state-of-the-art methods for sWGS analysis, widely used across the bioinformatics community; secondly, while allowing flexibility, it relieves the user of many data adjustment operations required for the interoperability between the different application, thus it is approachable even by entry-level bioinformaticians; thirdly, it heavily uses containerization (all the tools are offered through Docker or Singularity) to ensure that runs are as reproducible as possible; lastly, it builds upon a large *corpus* of existing, well-tested modules developed by the nf-core community [21].

Thus, SAMURAI represents a valuable first step towards standardized sWGS analysis. Future work will focus on incorporating additional copy number analysis methods and downstream analyses, further enhancing SAMURAI’s capabilities.

## Key Points

- SAMURAI offers a reproducible, scalable and standardized bioinformatics pipeline for the analysis of copy number alterations from shallow whole genome sequencing;
- SAMURAI’s copy number calling block offers two types of analysis workflow depending on the nature of the samples, one catering to the processing of tissues (“solid biopsy”) and another for the identification of aberrant CNAs in other biological fluids (“liquid biopsy”);
- SAMURAI provides a final report that allows an overall evaluation of the sequencing and alignment quality, as well as custom reports that facilitate the interpretation of copy number calling analysis.

## Funding

This work was supported by the Italian Association for Cancer Research (AIRC) [IG 29071 to C.R., IG 2024 - ID. 30381 to S.M.], EU funding within the MUR PNRR “National Center for HPC, BIG DATA AND QUANTUM COMPUTING” (Project no. CN00000013 CN1 to C.R.), the European Union’s Horizon 2020 research and innovation programme [grant agreement No 965193 for DECIDER to D.B and D.P.], and the Alessandra Bono Foundation to S.M.This research was supported by UniSMART, Fondazione Cassa di Risparmio di Padova e Rovigo, Intesa Sanpaolo, 1007 Confindustria Veneto Est, and AB ANALITICA S.r.l. under the initiative “Smart PhD”.

## Data availability

SAMURAI can be downloaded from https://github.com/DincalciLab/samurai. The additional scripts used for the analysis, along with configuration and expected outputs, are available at https://github.com/DIncalciLab/SAMURAI_paper_scripts. The real-world sequencing data sets used are deposited under controlled access at EGA (IDs EGAS00001004961 and EGAS00001004670).

## Acknowledgments

We would also like to thank the nf-core community for developing the nf-core infrastructure and resources for Nextflow pipelines. A full list of nf-core community members is available at https://nf-co.re/community.

## Authors’ contributions

**Conceptualization:** S.P., L.B., and C.R.

**Data curation:** S.P., L.B., and C.R.

**Formal analysis:** S.P. and L.B.

**Funding acquisition:** D.B., D.P., S.M., and C.R.

**Investigation:** S.P., L.B., and C.R.

**Methodology:** S.P., L.B., and C.R.

**Project administration:** D.B., D.P., S.M., and C.R.

**Resources:** D.B., D.P., L.B., and C.R.

**Software:** S.P., L.B., and C.R.

**Supervision:** D.B., D.P., S.M., L.B., and C.R.

**Validation:** L.B. and C.R.

**Visualization:** S.P.

**Writing - original draft:** S.P., L.B., and C.R.

**Writing - review & editing:** All authors

## Authors’ descriptions

**Sara Potente** is a PhD student at the department of Biology, University of Padova, Italy. She has been working on bioinformatics mainly applied to cancer genomics.

**Diego Boscarino** is the director of research projects co-funded through public grants at AB ANALITICA, developing new in vitro diagnostics devices based on molecular technologies, mainly Next Generation Sequencing.

**Dino Paladin** is the Founder of AB ANALITICA, Biofield Innovation, TEST VERITAS, DOTT DINO PALADIN company. Partner in Start-up: A MERAS ANNOS, DIGITALREHAB, OFFXET, EZ LAB. Inventor of 5 patents in diagnostic field and molecular biology. Responsible for several research projects.

**Sergio Marchini** is the head of Translational genomic Unit and the head of Humanitas Genomic Facility. He is in charge of different translational programs focused on monitoring minimal residual disease through liquid biopsy approach and improve early diagnosis of ovarian cancer

**Luca Beltrame** is a senior bioinformatician at IRCCS Humanitas Research Hospital, Rozzano, Milan, Italy. He is currently working in the study of genomic and epigenomic alterations in ovarian cancer, and on the development of reproducible workflows for bioinformatics analysis.

**Chiara Romualdi** is a full professor at the Department of Biology, University of Padova, Italy. She’s interested in the computational methods for genomic data analysis and integrations.

## Conflict of interest

The authors declare no competing financial interests.

## Notes

### Competing Interest Statement

The authors have declared no competing interest.

### Summary of Updates

Discussione section updated to clarify pipeline features and results interpretation; Some figures caption have been improved; supplementary files updated; Added an Aknowledgments section; References revised in order to be consistent between them.

