## Supplementary Table S1 for "SAMURAI: Shallow Analysis of copy nuMber alterations Using a Reproducible And Integrated bioinformatics pipeline"

| **Sample** | **Mean Depth** | **Median Depth** |
| --- | --- | --- |
| **SM-74_01** | **0.3110X** | **0X** |
| **SM-74_02** | **0.0548X** | **0X** |
| **SM-74_03** | **0.1866X** | **0X** |
| **SM-74_04** | **0.3110X** | **0X** |
| **SM-74_05** | **0.0548X** | **0X** |
| **SM-74_06** | **0.1866X** | **0X** |
| **SM-74_07** | **0.3110X** | **0X** |
| **SM-74_08** | **0.0548X** | **0X** |
| **SM-74_09** | **0.1866X** | **0X** |
| **SM-74_10** | **0.3110X** | **0X** |
| **SM-74_11** | **0.0548X** | **0X** |
| **SM-74_12** | **0.1866X** | **0X** |
| **SM-74_13** | **0.3110X** | **0X** |
| **SM-74_14** | **0.0548X** | **0X** |
| **SM-74_15** | **0.1866X** | **0X** |
| **SM-74_16** | **0.3110X** | **0X** |
| **SM-74_17** | **0.0548X** | **0X** |
| **SM-74_18** | **0.1866X** | **0X** |
| **SM-74_19** | **0.3110X** | **0X** |
| **SM-74_20** | **0.0548X** | **0X** |
| **SM-74_21** | **0.1866X** | **0X** |
| **SM-74_22** | **0.3110X** | **0X** |
| **SM-74_23** | **0.0548X** | **0X** |
| **SM-74_24** | **0.1866X** | **0X** |
| **SM-74_25** | **0.3110X** | **0X** |
| **SM-74_26** | **0.0548X** | **0X** |
| **SM-74_27** | **0.1866X** | **0X** |
| **SM-74_28** | **0.3110X** | **0X** |
| **SM-74_29** | **0.0548X** | **0X** |
| **SM-74_30** | **0.1866X** | **0X** |

**Supplementary Table S2. Mean and median coverage for *in-silico* dilutions of synthetic normal data (see Materials and Methods).**
