## Supplementary Figure S1 for "SAMURAI: Shallow Analysis of copy nuMber alterations Using a Reproducible And Integrated bioinformatics pipeline"

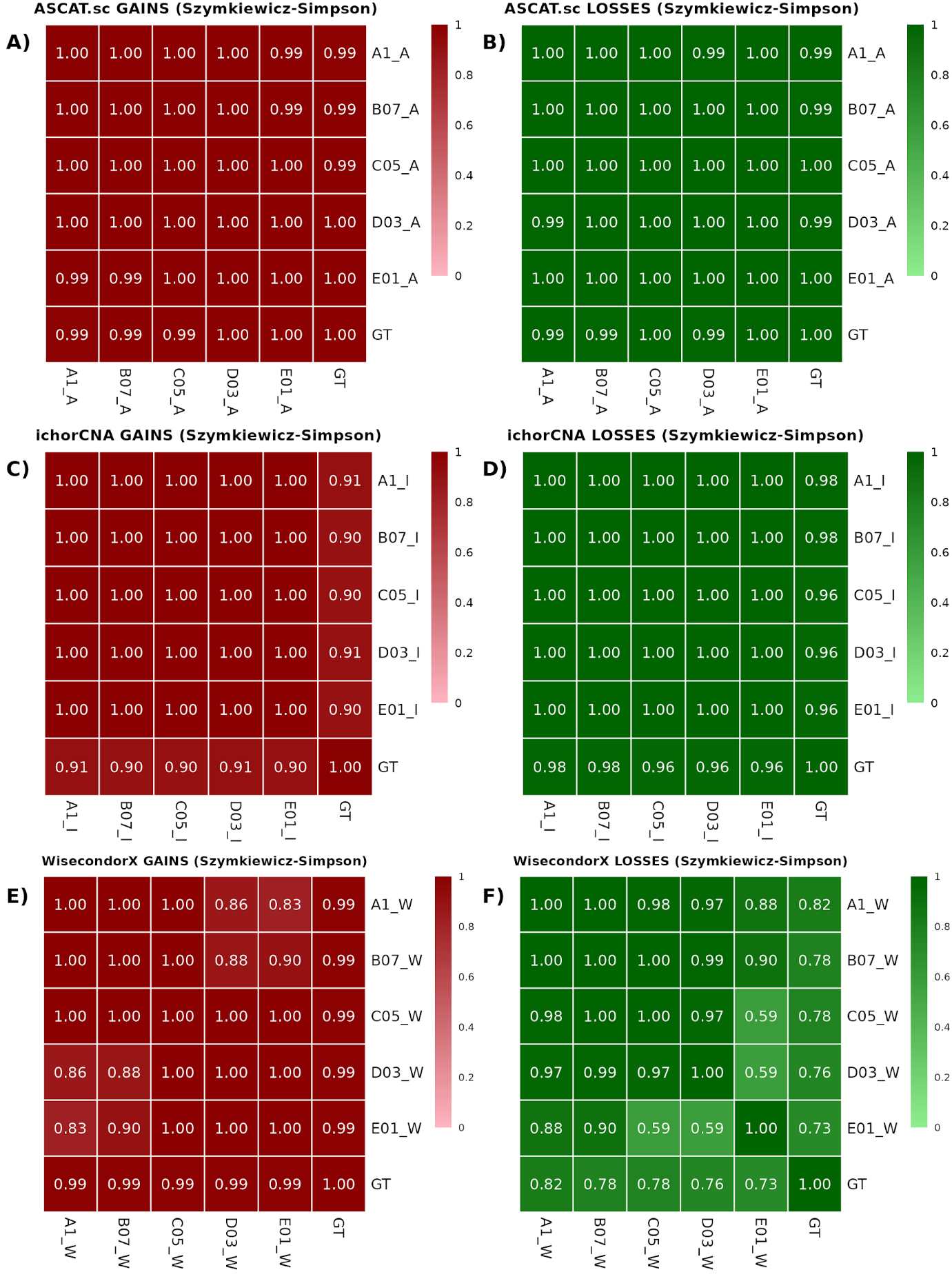

**Supplementary Figure S1**. Heatmap reporting the Szymkiewicz-Simpson coefficient metrics for the comparison between inferred copy numbers and ground truth copy numbers. A) Metrics for gains inferred from ASCAT.sc versus ground truth. B) Metrics for losses inferred from ASCAT.sc versus ground truth. C) Metrics for gains inferred from ichorCNA versus ground truth. D) Metrics for losses inferred from ichorCNA versus ground truth. E) Metrics for gains inferred from Wisecondorx versus ground truth. F) Metrics for losses inferred from WisecondorX versus ground truth.
Original sample: A1, *in-silico* diluted samples: B07, C05, D03, E01. Color intensity within the heatmap reflects the coefficient value, with higher values indicating better agreement.
