## Supplementary Table S3 for "SAMURAI: Shallow Analysis of copy nuMber alterations Using a Reproducible And Integrated bioinformatics pipeline"

| **HGSOC** | | |
| --- | --- | --- |
| **Sample ID** | **GAINS** | **LOSSES** |
| 1 | 0,97 | 0,99 |
| 10 | 0,92 | 1,00 |
| 113_left | 0,97 | 0,96 |
| 113_right | 0,99 | 0,95 |
| 119 | 0,88 | 0,97 |
| 12 | 0,99 | 1,00 |
| 123_left | 0,97 | 0,96 |
| 123_right | 0,93 | 0,73 |
| 131_left | 0,81 | 0,64 |
| 131_right | 0,91 | 0,97 |
| 133 | 0,97 | 0,99 |
| 137 | 1,00 | 0,96 |
| 148 | 1,00 | 0,98 |
| 164 | 0,98 | 0,98 |
| 183_left | 0,86 | 0,82 |
| 183_right | 0,97 | 0,47 |
| 186 | 0,99 | 0,08 |
| 202_left | 1,00 | 0,99 |
| 202_right | 0,98 | 0,99 |
| 203 | 0,99 | 0,98 |
| 206 | None (SAMURAI) | 0,00 |
| 207 | 0,96 | 0,62 |
| 21 | 0,97 | 0,99 |
| 212 | 1,00 | 0,99 |
| 219 | 0,99 | 0,91 |
| 31 | 0,91 | 0,69 |
| 31bis | 0,98 | 0,68 |
| 31tris | 0,93 | 0,98 |
| 4 | 0,98 | 0,99 |
| 41 | 0,91 | 0,99 |
| 45 | 0,98 | 0,00 |
| 5 | 0,95 | 0,95 |
| 50 | 1,00 | 0,96 |
| 51 | 0,98 | 0,06 |
| 8 | 0,97 | 1,00 |
| 89 | 0,92 | 1,00 |
| 99 | 1,00 | 0,99 |
| T15 | 1,00 | 0,99 |
| T20 | 0,97 | 0,99 |
| T25 | 0,90 | 0,79 |
| T37 | 0,98 | 1,00 |
| T41 | 0,94 | 0,99 |
| T5 | 0,99 | None (SAMURAI) |
| T59 | 0,16 | 0,36 |
| T6 | 0,95 | 0,98 |
| T62 | 0,99 | 0,76 |
| T9 | 0,82 | 0,98 |
| **LGSOC** | | |
| **Sample ID** | **GAINS** | **LOSSES** |
| 106 | 1,00 | 1,00 |
| 13 | 0,99 | 1,00 |
| 151 | 1,00 | 1,00 |
| 156 | 1,00 | None |
| 161_left | 0,00 | 0,00 |
| 161_right | 0,00 | 0,20 |
| 175 | 0,79 | 0,99 |
| 18 | None (SAMURAI) | 0,00 |
| 187_left | 1,00 | 0,98 |
| 187_right | 1,00 | 0,99 |
| 20 | 0,97 | 0,89 |
| 205 | 0,99 | 0,98 |
| 215 | 0,94 | None (Data set T) |
| 26 | None (SAMURAI) | None (Data set T) |
| 3 | 0,92 | 0,96 |
| 39 | 1,00 | 0,99 |
| 52 | 1,00 | 1,00 |
| 75 | 0,68 | 0,14 |
| T27 | 1,00 | 0,02 |
| T32 | 0,00 | 0,03 |
| **EC** | | |
| **Sample ID** | **GAINS** | **LOSSES** |
| 100 | 0,92 | 0,08 |
| 101 | 1,00 | 0,61 |
| 104 | 0,92 | 0,98 |
| 105 | None | None (Data set T) |
| 107 | 1,00 | 0,87 |
| 11 | 0,88 | 0,00 |
| 111 | 0,00 | 0,65 |
| 112 | 0,74 | 0,00 |
| 114 | 1,00 | 1,00 |
| 116 | 0,60 | 0,03 |
| 117 | 0,17 | 0,26 |
| 127 | 0,95 | 0,82 |
| 129 | None (Data set T) | 0,25 |
| 134 | 0,98 | None (Data set T) |
| 147 | 0,98 | 0,89 |
| 149 | 0,86 | 0,97 |
| 150 | 0,43 | 1,00 |
| 154 | 0,97 | 0,83 |
| 158 | 0,84 | 0,96 |
| 159 | 1,00 | 1,00 |
| 163 | 0,86 | 0,65 |
| 17 | None (Data set T) | 0,00 |
| 178 | 1,00 | 0,73 |
| 179 | 1,00 | 0,75 |
| 180 | 0,40 | 0,09 |
| 182 | 1,00 | 1,00 |
| 184 | 0,97 | 0,99 |
| 185 | 0,00 | 0,04 |
| 188 | 1,00 | 1,00 |
| 188bis | 0,94 | 0,98 |
| 190 | 0,84 | 0,76 |
| 191 | 0,00 | 0,00 |
| 192 | 1,00 | 0,97 |
| 195 | 1,00 | 0,90 |
| 198 | 1,00 | 1,00 |
| 200 | 0,95 | None (SAMURAI) |
| 210 | 0,94 | 0,98 |
| 22 | 0,96 | 0,08 |
| 221 | 0,92 | 0,94 |
| 223 | 0,95 | 0,99 |
| 225 | 1,00 | 1,00 |
| 28 | 0,00 | 0,47 |
| 37 | None (Data set T) | None (Data set T) |
| 38 | 1,00 | None (Data set T) |
| 42 | None (Data set T) | 0,97 |
| 44 | 0,00 | None |
| 47 | 1,00 | None (Data set T) |
| 49 | 0,89 | 0,93 |
| 54 | 0,96 | 0,99 |
| 59 | 0,00 | 0,63 |
| 60 | 0,85 | 0,05 |
| 65 | 1,00 | 1,00 |
| 72 | 0,99 | 0,79 |
| 73 | 0,99 | 1,00 |
| 80 | 0,99 | 0,99 |
| 84 | 1,00 | None (Data set T) |
| 85 | 1,00 | 0,18 |
| 90 | 0,99 | 0,94 |
| 91 | 0,98 | 0,98 |
| 92 | 1,00 | None |
| 96 | 0,73 | 0,65 |
| 97 | 0,99 | 0,78 |
| 97bis | 1,00 | 0,00 |
| 98 | 1,00 | 0,86 |
| T10 | 0,73 | 0,63 |
| T16 | 0,00 | 0,00 |
| T2 | None (SAMURAI) | 0,13 |
| T22 | 0,97 | 0,95 |
| T23 | 0,69 | 1,00 |
| T24 | 0,97 | 0,00 |
| T28 | None (SAMURAI) | 0,38 |
| T29 | 0,00 | 0,00 |
| T3 | 0,99 | 0,29 |
| T33 | 1,00 | 0,97 |
| T4 | 1,00 | 0,99 |
| T42 | 0,95 | 0,99 |
| T47 | 0,96 | 0,99 |
| T48 | 0,99 | 0,00 |
| T52 | 1,00 | 0,96 |
| T55 | 0,61 | 0,97 |
| T57 | 0,98 | 0,69 |
| T8 | 0,97 | 0,98 |
| **MOC** | | |
| **Sample ID** | **GAINS** | **LOSSES** |
| 102 | 0,61 | 0,08 |
| 103 | None (Data set T) | 0,95 |
| 118 | 1,00 | 1,00 |
| 122 | 0,70 | 0,98 |
| 124 | 0,91 | 0,98 |
| 135 | None | None (Data set T) |
| 14 | 0,73 | 0,88 |
| 141 | None (SAMURAI) | 0,17 |
| 15 | 0,93 | 0,72 |
| 155 | 1,00 | 0,99 |
| 166 | None (SAMURAI) | 0,99 |
| 168 | 0,99 | 0,98 |
| 171 | 0,64 | 0,91 |
| 172 | 0,99 | 1,00 |
| 176 | 0,93 | 0,00 |
| 189 | 0,00 | 0,99 |
| 197 | None (SAMURAI) | None (Data set T) |
| 204 | None (Data set T) | 0,00 |
| 214 | 0,84 | 0,98 |
| 217 | 1,00 | 0,00 |
| 25 | 0,00 | None (Data set T) |
| 34 | 0,96 | 0,90 |
| 36 | 0,98 | 1,00 |
| 48 | 1,00 | 1,00 |
| 53 | 1,00 | 1,00 |
| 57 | 0,82 | 0,25 |
| 63 | 0,94 | 0,95 |
| 67 | 1,00 | None (Data set T) |
| 69 | None (SAMURAI) | 0,99 |
| 7 | 0,95 | 0,00 |
| T14 | 0,87 | 1,00 |
| T26 | 0,32 | 0,82 |
| T31 | 0,98 | 0,93 |
| T38 | 0,98 | 0,92 |
| T39 | 0,75 | 1,00 |
| T44 | 1,00 | 0,62 |
| T53 | 0,99 | 0,91 |
| T60 | 0,00 | 0,53 |
| **OCCC** | | |
| **Sample ID** | **GAINS** | **LOSSES** |
| 115 | 0,71 | 0,12 |
| 115bis | 1,00 | None (Data set T) |
| 126 | 0,99 | 1,00 |
| 128 | 0,98 | 0,82 |
| 132 | 0,13 | 0,27 |
| 16 | 0,97 | 0,98 |
| 174 | 0,81 | 0,98 |
| 181 | 0,94 | 0,97 |
| 193 | 0,98 | 0,00 |
| 208 | 0,99 | 0,98 |
| 213 | 0,98 | 0,98 |
| 220 | 0,95 | 0,97 |
| 224 | 0,94 | 0,95 |
| 23_left | 0,65 | 0,49 |
| 23_right | 1,00 | 0,78 |
| 29 | 0,98 | 0,99 |
| 33 | 0,97 | 1,00 |
| 6 | 0,75 | 0,79 |
| 70 | 1,00 | 0,99 |
| 78 | 0,97 | 0,95 |
| 82 | 1,00 | 1,00 |
| 83 | None (SAMURAI) | 1,00 |
| 86 | 1,00 | 0,98 |
| 95 | 0,88 | 0,98 |
| T1 | 0,98 | 0,98 |
| T13 | 1,00 | 0,96 |
| T19 | 0,00 | 0,53 |
| T36 | 0,98 | 0,75 |
| T49 | 0,98 | 0,99 |
| T54 | 1,00 | 0,98 |
| T56 | 1,00 | 0,16 |

**Supplementary Table S3.** Overlap coefficient between data in data set T from original work and SAMURAI results for each sample in the cohort, divided by histotype. None indicates that no copy number alterations were found in both the original data and the SAMURAI analysis and thus a coefficient was not calculated; None (SAMURAI) indicates that it was due to no CNAs found by SAMURAI, and None (Data set T) due to no CNAs present in the original data set.
