## Supplementary Figure S2 for "SAMURAI: Shallow Analysis of copy nuMber alterations Using a Reproducible And Integrated bioinformatics pipeline"

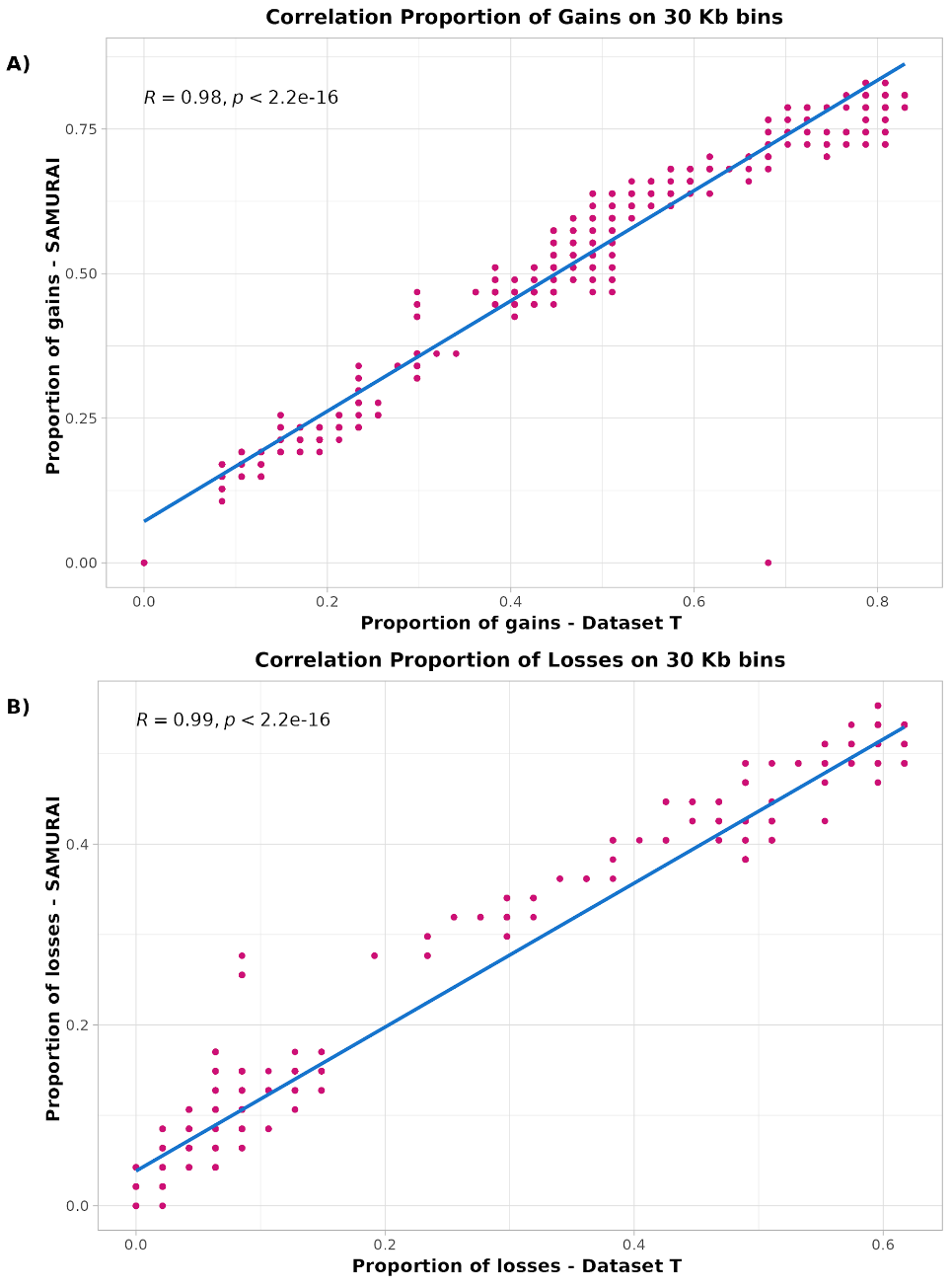

**Supplementary Figure S2.** Scatterplot showing the correlation between proportion of gains and losses on overlapping 30 Kbp bins of chr8 between original segmentation files and SAMURAI’s segmentation files of HGSOC samples in data set T. **A)** Correlation between proportions of copy number gains. x axis, proportions from computed from original data; y axis, proportions from SAMURAI data. **B)** Correlation between proportions of copy number losses. x axis, proportions from computed from original data; y axis, proportions from SAMURAI data. R: Pearson Correlation coefficient; *p*: p-value.
