## Supplementary Figure S3 for "SAMURAI: Shallow Analysis of copy nuMber alterations Using a Reproducible And Integrated bioinformatics pipeline"

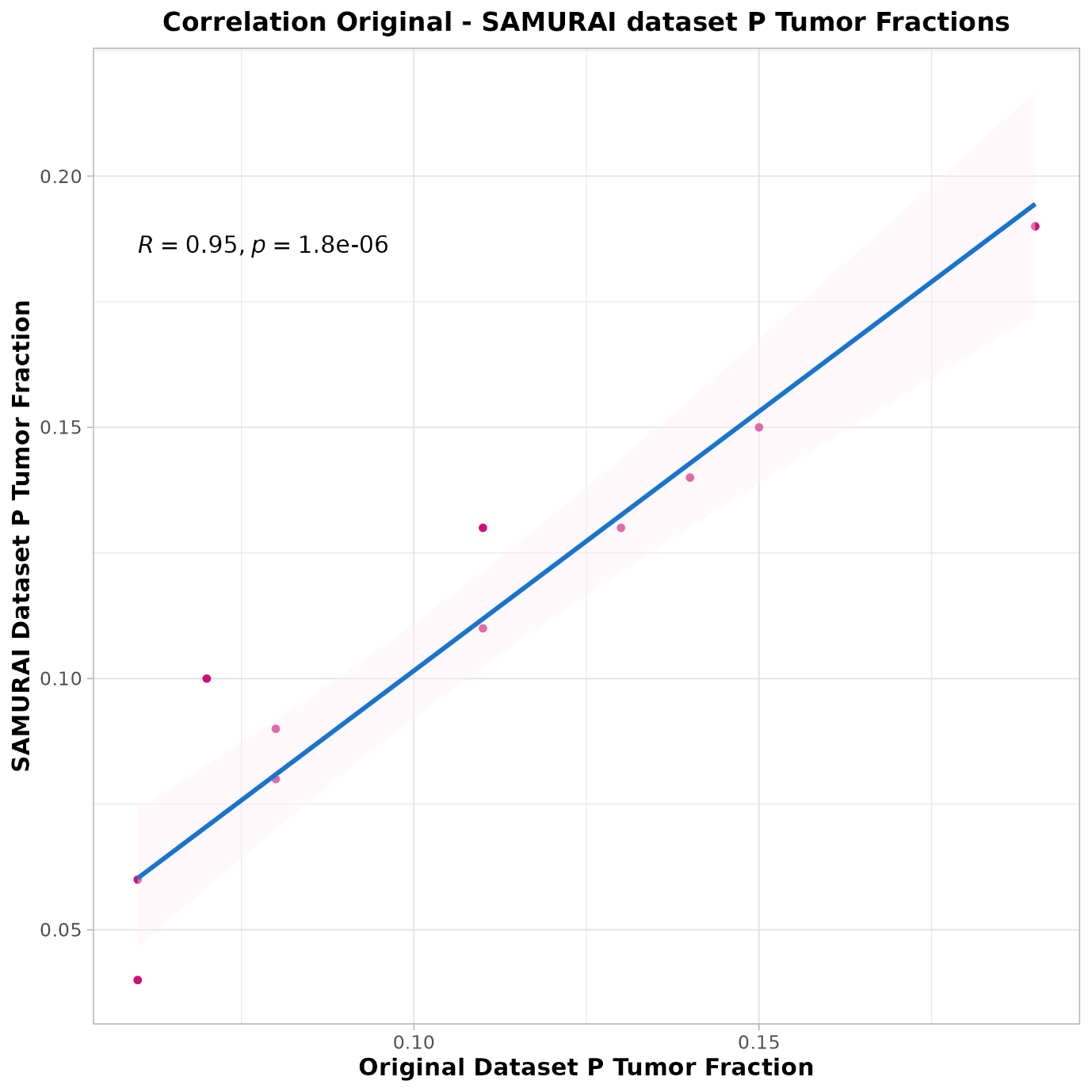

**Supplementary Figure S3.** Scatterplot illustrating the correlation between tumor fractions (TFs) computed by SAMURAI with ichorCNA workflow and the original reported tumor fractions in data set P. X axis, original TFs from data set P; y axis, TFs computed by SAMURAI. R: Pearson Correlation coefficient; *p*: p-value.
