## Supplementary File 1 for "SAMURAI: Shallow Analysis of copy nuMber alterations Using a Reproducible And Integrated bioinformatics pipeline"

multiqc v1.21
SAMURAI Results
- General Stats
- Picard
  - Alignment Summary
  - Mean read length
  - Base Distribution
  - Insert Size
  - Mean Base Quality by Cycle
  - Base Quality Distribution
  - WGS Coverage
  - WGS Filtered Bases
- Gistic arm-level analysis summary
- Gistic Analysis
- ASCAT.sc Summary
- Gistic Analysis Summary
- Copy Number Signatures Activities
- Software Versions
- dincalcilab/samurai Methods Description
- dincalcilab/samurai Workflow Summary


multiqc

### SAMURAI Results

SAMURAI's analysis results of Copy Number Alterations from shallow whole genome sequencing.

A modular tool to aggregate results from bioinformatics analyses across many samples into a single report.

This report has been generated by the dincalcilab/samurai analysis pipeline.

Report generated on 2024-11-28, 16:47 CET based on data in: `/home/incalci/lbeltrame/analysis/samurai-stage2/work/e2/3e1d51567d1684a41e79e9c3351757`

Welcome! Not sure where to start? Watch a tutorial video (6:06) don't show again

#### General Statistics

Copy table | Configure columns | Scatter plot | Violin plot | Showing 46/46 rows and 2/6 columns. | Export as CSV

| Sample Name | Insert Size | Mean Coverage |
|---|---|---|
| 1 | 221 bp | 0.2 X |
| 10 | 242 bp | 0.3 X |
| 113_left | 105 bp | 0.2 X |
| 113_right | 105 bp | 0.2 X |
| 119 | 197 bp | 0.2 X |
| 12 | 210 bp | 0.3 X |
| 123_left | 211 bp | 0.4 X |
| 123_right | 212 bp | 0.4 X |
| 131_left | 209 bp | 0.2 X |
| 131_right | 224 bp | 0.3 X |
| 133 | 221 bp | 0.4 X |
| 137 | 194 bp | 0.3 X |
| 148 | 240 bp | 0.4 X |
| 164 | 234 bp | 0.3 X |
| 183_left | 100 bp | 0.2 X |

#### Picard

Picard is a set of Java command line tools for manipulating high-throughput sequencing data.

##### Alignment Summary

Please note that Picard's read counts are divided by two for paired-end data. Total bases (including unaligned) is not provided.

Percentages | Aligned Reads | Aligned Bases | Export Plot

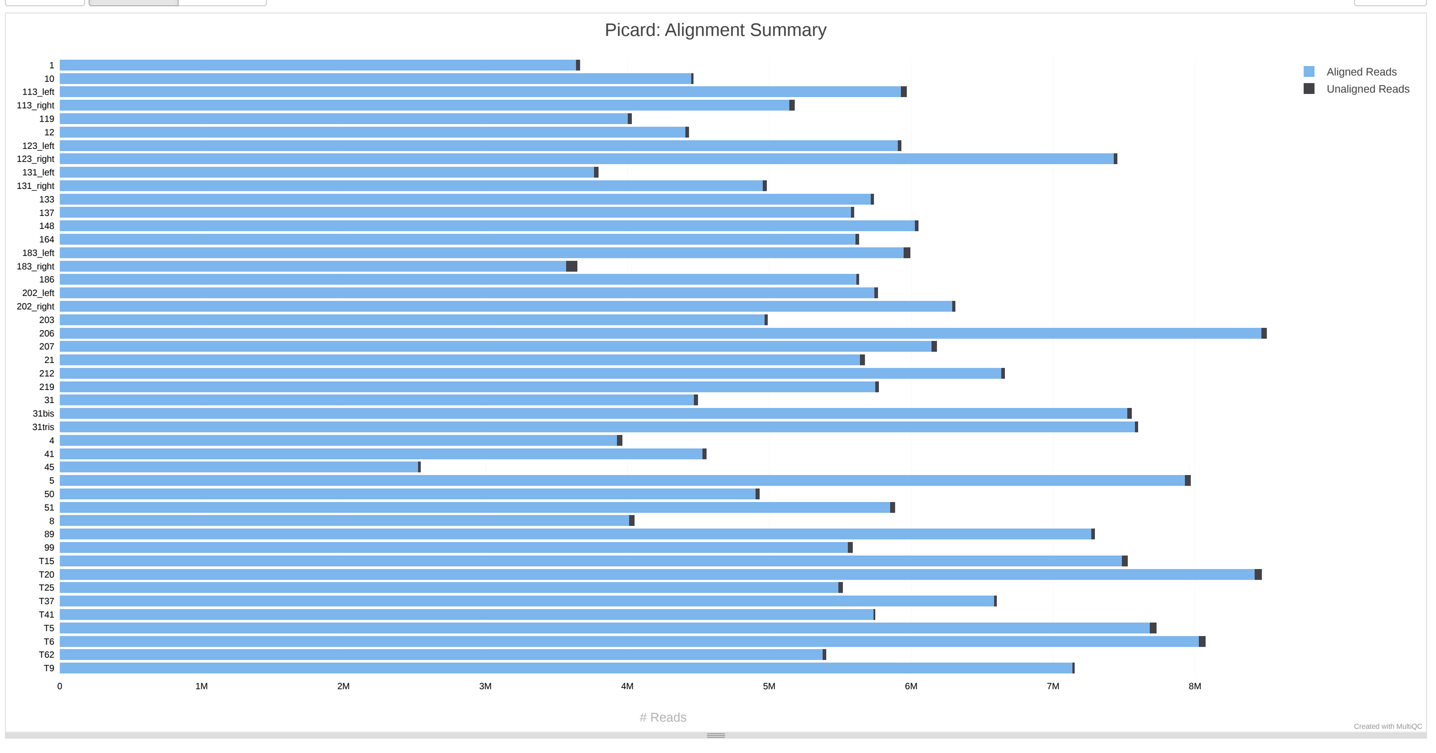


##### Mean read length

The mean read length of the set of reads examined.

Export Plot

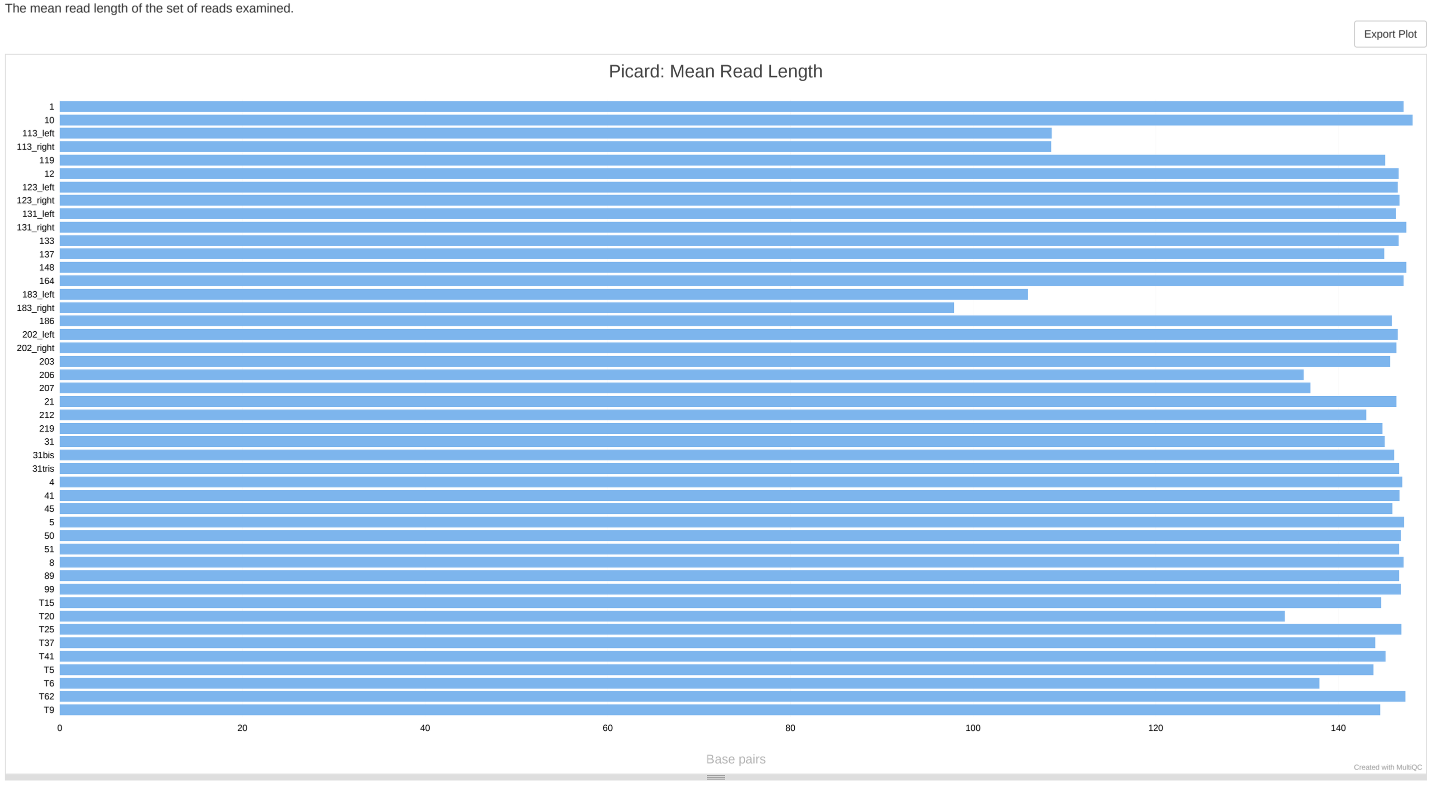


##### Base Distribution

Plot shows the distribution of bases by cycle.

% Adenine | % Cytosine | % Guanine | % Thymine | % Undetermined | Export Plot

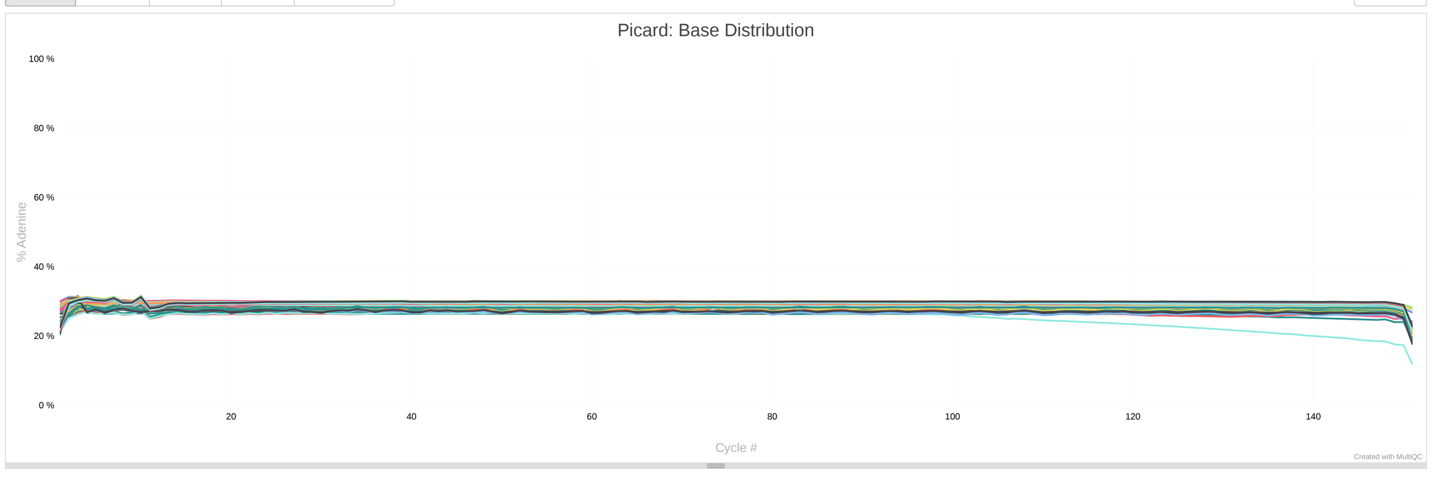


##### Insert Size

Plot shows the number of reads at a given insert size. Reads with different orientations are summed.

Counts | Percentages | Export Plot

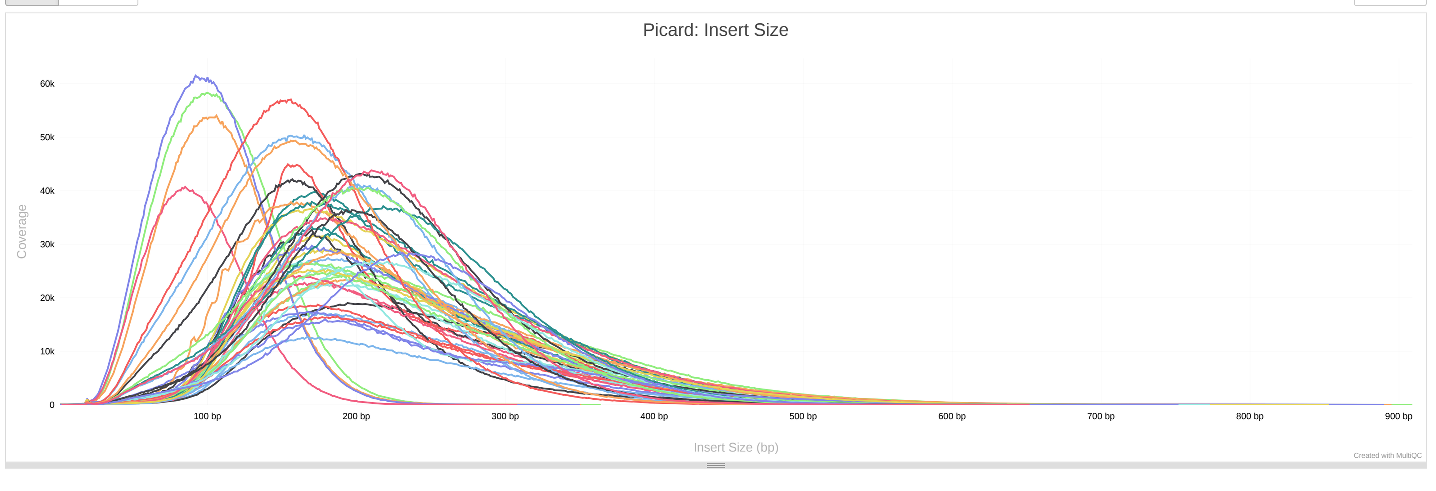


##### Mean Base Quality by Cycle

Plot shows the mean base quality by cycle.

Help | Export Plot

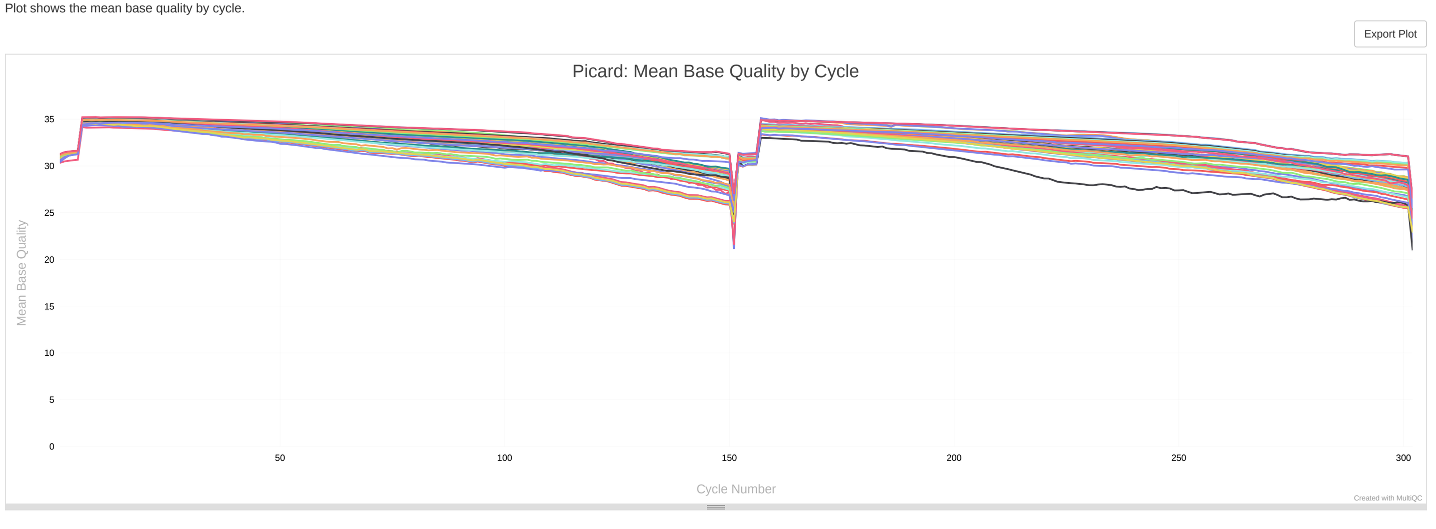


##### Base Quality Distribution

Plot shows the count of each base quality score.

Export Plot

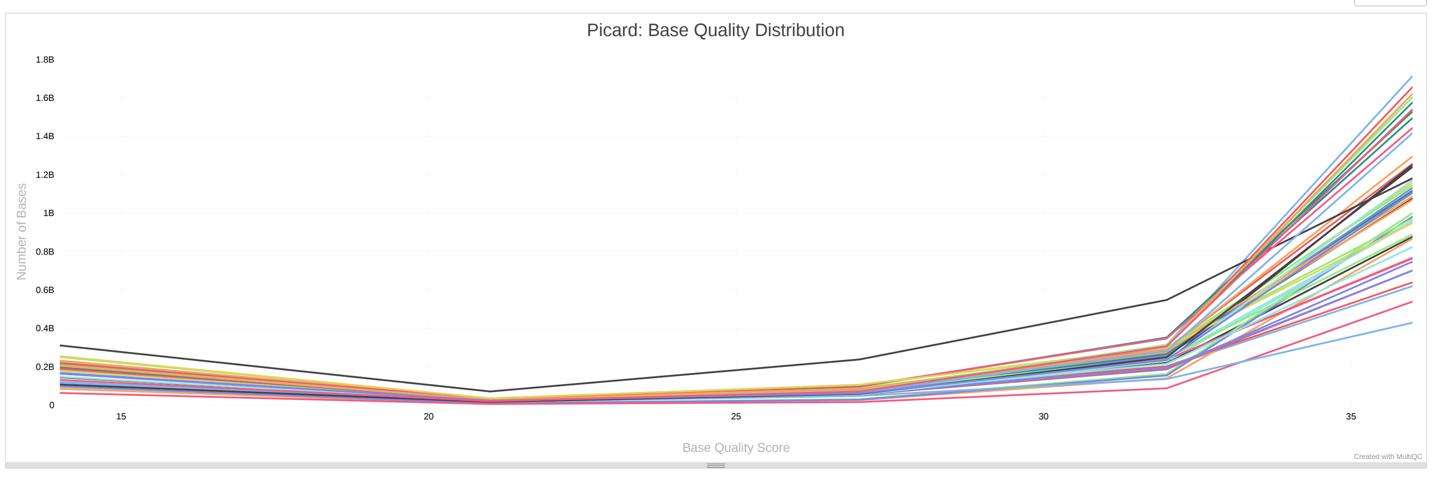


##### WGS Coverage

The number of bases in the genome territory for each fold coverage. Note that final 1% of data is hidden to prevent very long tails.

Percentage Drop-Off | Counts Histogram | Export Plot

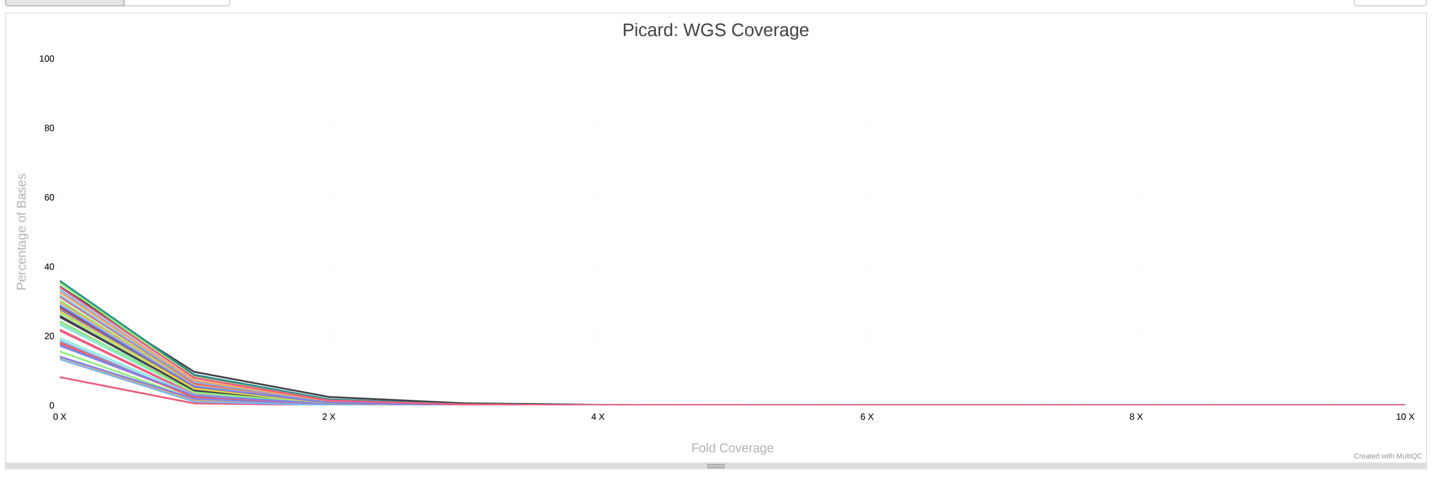


##### WGS Filtered Bases

For more information about the filtered categories, see the Picard documentation.

Export Plot

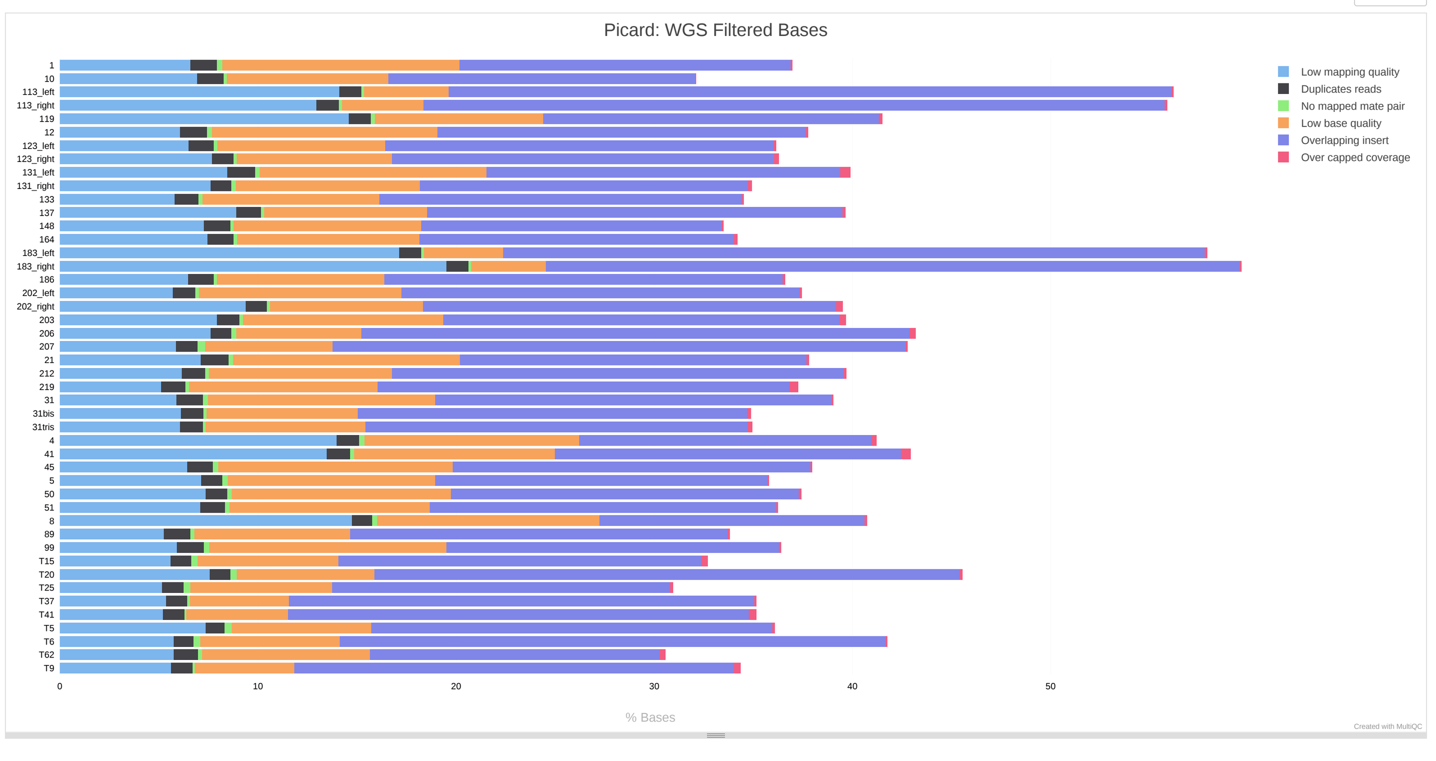


#### Gistic arm-level analysis summary

This table shows significant recurrent arm-level alterations identified by GISTIC2. Each alteration has an associated frequency and q-value for statistical significance. .

Copy table | Configure columns | Scatter plot | Violin plot | Showing 10/10 rows and 4/4 columns. | Export as CSV

| Arm | Type of alteration | No. of samples | Frequency | q-value |
|---|---|---|---|---|
| 4p | Del | 20 | 43.0 | 0.0145 |
| 4q | Del | 21 | 46.0 | 0.0011 |
| 8p | Del | 24 | 52.0 | 0.0001 |
| 13q | Del | 22 | 48.0 | 0.0012 |
| 16p | Del | 18 | 39.0 | 0.0158 |
| 16q | Del | 20 | 43.0 | 0.0051 |
| 17p | Del | 33 | 70.0 | 0.0000 |
| 17q | Del | 25 | 52.0 | 0.0000 |
| 18q | Del | 19 | 41.0 | 0.0344 |
| 22q | Del | 21 | 43.0 | 0.0097 |

#### Gistic Analysis

Maftools genomic plot with segments highlighting significant Amplifications and Deletions.

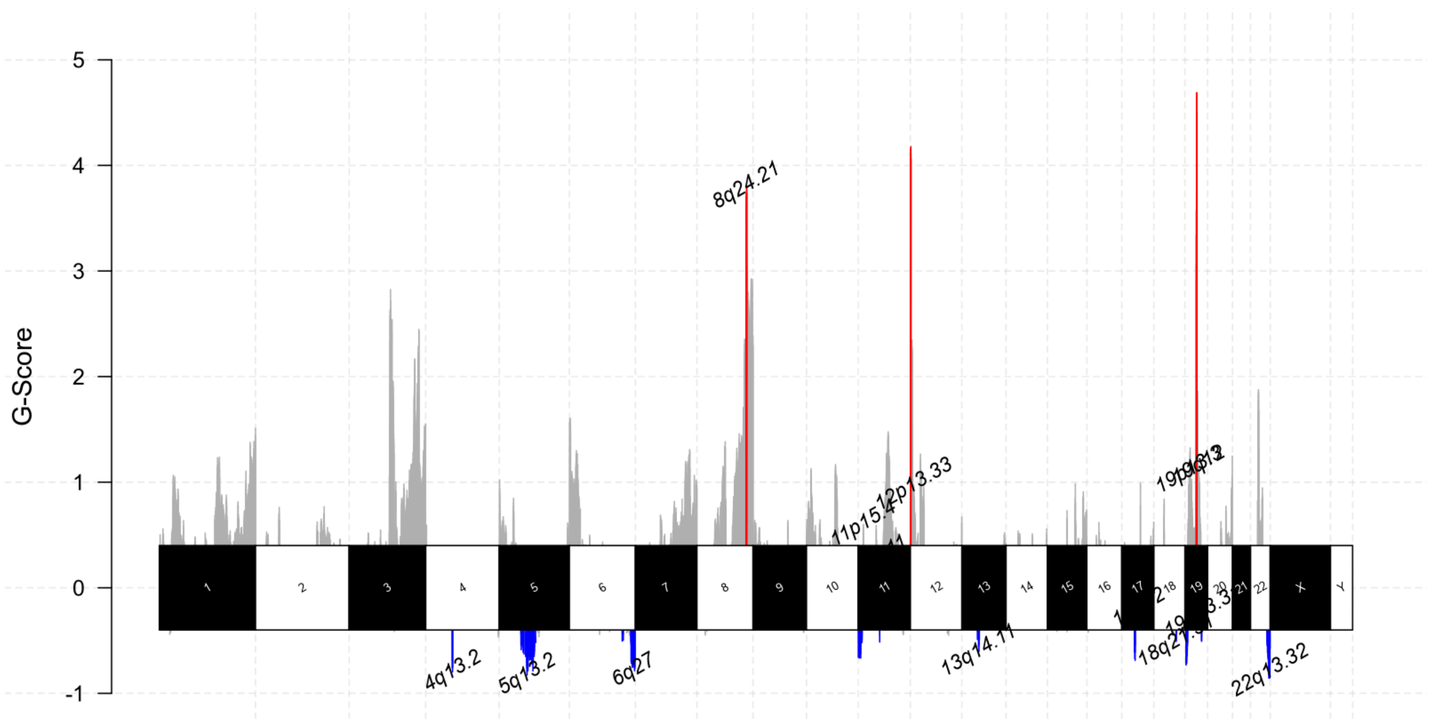


#### ASCAT.sc Summary

This table shows purity and ploidy values computed by ASCAT.sc.

Copy table | Configure columns | Scatter plot | Violin plot | Showing 46/46 rows and 2/2 columns. | Export as CSV

| Sample ID | Purity | Ploidy |
|---|---|---|
| T37 | 0.4700 | 2.4200 |
| 186.0 | 0.2900 | 2.7600 |
| T5 | 1.0000 | 2.0100 |
| 1.0 | 0.8600 | 2.8200 |
| 99.0 | 0.9600 | 3.8600 |
| 5.0 | 0.7800 | 2.8200 |
| T41 | 0.7700 | 1.7900 |
| 148.0 | 0.9200 | 2.9200 |
| 31_Tris | 0.9000 | 4.4900 |
| 164.0 | 0.9000 | 1.8000 |
| T25 | 0.7400 | 3.1400 |
| 51.0 | 0.5000 | 3.4200 |
| 131_right | 0.6400 | 2.7700 |
| 50.0 | 0.7200 | 3.0900 |
| 113_right | 0.8300 | 1.7500 |

#### Gistic Analysis Summary

This table shows significant lesions identified by GISTIC2 analysis. Focal lesions are alterations which span less than 25% of a chromosomal arm. Broad lesions span more than 25% of the chromosomal arm, but less than a whole arm .

Copy table | Configure columns | Scatter plot | Violin plot | Showing 14/14 rows and 9/9 columns. | Export as CSV

| Chromosome | Chromosome | Cytoband | Start | End | Length | CN Kind | Lesion Frequency | Nr. of Samples | Type of Lesion |
|---|---|---|---|---|---|---|---|---|---|
| Del_4q13.2_chr4:68120002-69420000 | chr4 | 4q13.2 | 68 120 002 | 69 420 000 | 1 299 998 | Del | 0.50 | 23 | focal |
| Del_5q13.2_chr5:55290627-94745000 | chr5 | 5q13.2 | 55 290 627 | 94 745 000 | 39 454 373 | Del | 0.59 | 27 | focal |
| Del_6q27_chr6:159125002-170805979 | chr6 | 6q27 | 159 125 002 | 170 805 979 | 11 680 977 | Del | 0.70 | 32 | focal |
| Amp_8q24.21_chr8:127190558-129010000 | chr8 | 8q24.21 | 127 190 558 | 129 010 000 | 1 819 442 | Amp | 0.80 | 37 | focal |
| Del_11p15.4_chr11:1-10565000 | chr11 | 11p15.4 | 1 | 10 565 000 | 10 564 999 | Del | 0.65 | 30 | focal |
| Del_11q11_chr11:55370002-55710000 | chr11 | 11q11 | 55 370 002 | 55 710 000 | 339 998 | Del | 0.33 | 15 | focal |
| Amp_12p13.33_chr12:760002-1680000 | chr12 | 12p13.33 | 760 002 | 1 680 000 | 919 998 | Amp | 0.63 | 29 | focal |
| Del_13q14.11_chr13:31620387-54774926 | chr13 | 13q14.11 | 31 620 387 | 54 774 926 | 23 154 539 | Del | 0.57 | 26 | focal |
| Del_17q12_chr17:32765002-34935000 | chr17 | 17q12 | 32 765 002 | 34 935 000 | 2 169 998 | Del | 0.70 | 32 | focal |
| Del_18q21.31_chr18:37155063-80373285 | chr18 | 18q21.31 | 37 155 063 | 80 373 285 | 43 218 222 | Del | 0.61 | 28 | broad |
| Amp_19q12_chr19:29950002-30550000 | chr19 | 19q12 | 29 950 002 | 30 550 000 | 599 998 | Amp | 0.43 | 20 | focal |
| Del_19p13.3_chr19:1-6220000 | chr19 | 19p13.3 | 1 | 6 220 000 | 6 219 999 | Del | 0.63 | 29 | broad |
| Del_19q13.31_chr19:42740002-43435000 | chr19 | 19q13.31 | 42 740 002 | 43 435 000 | 694 998 | Del | 0.48 | 22 | focal |
| Del_22q13.32_chr22:44505002-50204868 | chr22 | 22q13.32 | 44 505 002 | 50 204 868 | 5 699 866 | Del | 0.74 | 34 | focal |

#### Copy Number Signatures Activities

The image is a summary view of Copy number Signatures from segmentation files, computed by the R Package CINSignatureQuantification.

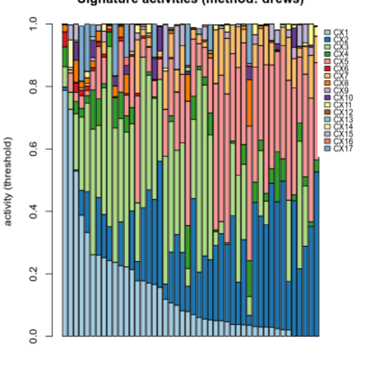


#### Software Versions

Software Versions lists versions of software tools extracted from file contents.

Copy table

| Group | Software | Version |
|---|---|---|
| ASCAT_SC | ASCAT.sc | 0.1 |
| ASSEMBLE_GISTIC_OUTPUT | gistic-cli | 0.4.5 |
| GISTIC2 | GISTIC2 | 0.1 |
| MAFTOOLS | MAFTOOLS | 0.1 |
| PICARD_COLLECTMULTIPLEMETRICS | picard | 3.1.1 |
| Workflow | Nextflow | 24.4.2 |
| | dincalcilab/samurai | v1.0.3-gdf6e344 |

#### dincalcilab/samurai Methods Description

Suggested text and references to use when describing pipeline usage within the methods section of a publication.

##### Methods

Data was processed using dincalcilab/samurai v1.0.3 (doi: 10.1101/2024.09.30.615766) of the nf-core collection of workflows (Ewels *et al.*, 2020).

The pipeline was executed with Nextflow v24.04.2 (Di Tommaso *et al.*, 2017) with the following command:

```
nextflow run DIncalciLab/swgscna -r master --input samplesheet.hgsoc.csv -params-file parameters.yaml --outdir output/ -profile cluster -resume -qs 2
```


Notes:

- The command above does not include parameters contained in any configs or profiles that may have been used. Ensure the config file is also uploaded with your publication!
- You should also cite all software used within this run. Check the "Software Versions" of this report to get version information.

#### dincalcilab/samurai Workflow Summary

- this information is collected when the pipeline is started.

**Core Nextflow options**

| | |
|---|---|
| revision | master |
| runName | evil_babbage |
| containerEngine | apptainer |
| launchDir | /home/incalci/lbeltrame/analysis/samurai-stage2 |
| workDir | /home/incalci/lbeltrame/analysis/samurai-stage2/work |
| projectDir | /home/incalci/lbeltrame/.nextflow/assets/DIncalciLab/swgscna |
| userName | lbeltrame |
| profile | cluster |
| configFiles | N/A |

**Input/output options**

| | |
|---|---|
| input | samplesheet.hgsoc.csv |
| outdir | output/ |

**Reference genome options**

| | |
|---|---|
| fasta | /opt/bcbio/1.2.8/genomes/Hsapiens/hg38/seq/hg38.fa |
| fai | /opt/bcbio/1.2.8/genomes/Hsapiens/hg38/seq/hg38.fa.fai |
| dict | /opt/bcbio/1.2.8/genomes/Hsapiens/hg38/seq/hg38.dict |
| igenomes_ignore | true |

**Institutional config options**

| | |
|---|---|
| custom_config_base | /home/incalci/shared/pipeline-configs/ |

**Max job request options**

| | |
|---|---|
| max_cpus | 100 |
| max_memory | 260.GB |

**Generic options**

| | |
|---|---|
| hook_url | https://hookshot.heavensinferno.net/webhook/slack/a154b5f4-5a45-491c-9d4c-fffa71e7dd4d |
| multiqc_logo | /home/incalci/lbeltrame/.nextflow/assets/DIncalciLab/swgscna/assets/dincalcilab-samurai_logo_dark.png |

**Common sWGS options**

| | |
|---|---|
| binsize | 30 |
| caller | ascat_sc |

**ichorCNA options**

| | |
|---|---|
| ichorcna_chrs_to_use | paste0('chr', c(1:22)) |
| ichorcna_chrs_to_train | paste0('chr', c(1:22)) |
| ichorcna_chrs_to_no… | paste0('chr', c(1:22)) |

**GISTIC specific options**

| | |
|---|---|
| run_gistic | true |
| gistic_t_amp | 0.3 |
| gistic_t_del | 0.3 |
| gistic_remove_x | true |
| gistic_broad_analysis | true |
| gistic_broad_chr_len… | 0.7 |

**Other parameters**

| | |
|---|---|
| compute_signatures | true |
