## Supplementary File 2 for "SAMURAI: Shallow Analysis of copy nuMber alterations Using a Reproducible And Integrated bioinformatics pipeline"

multiqc v1.21
SAMURAI Results
General Stats
Picard
Alignment Summary
Mean read length
Base Distribution
Insert Size
Mean Base Quality by Cycle
Base Quality Distribution
WGS Coverage
WGS Filtered Bases
Gistic arm-level analysis summary
IchorCNA
Gistic Analysis
Gistic Analysis Summary
Software Versions
dincalcilab/samurai Methods Description
dincalcilab/samurai Workflow Summary

multiqc

### SAMURAI Results

SAMURAI's analysis results of Copy Number Alterations from shallow whole genome sequencing.

A modular tool to aggregate results from bioinformatics analyses across many samples into a single report.

This report has been generated by the dincalcilab/samurai analysis pipeline.

Report generated on 2024-07-26, 15:24 CEST based on data in: `/home/incalci/lbeltrame/analysis/samurai-ichorcna/work/71/ce24b3ae34d027a022f66676b5e1a2`

Welcome! Not sure where to start? Watch a tutorial video (6:06) don't show again

#### General Statistics

Copy table | Configure columns | Scatter plot | Violin plot | Showing 12/12 rows and 2/6 columns. | Export as CSV

| Sample Name | Insert Size | Mean Coverage |
|---|---|---|
| 21531-PL1_cfdna_ready | 147 bp | 0.5 X |
| 21553-PL1_cfdna_ready | 173 bp | 0.3 X |
| 21557-PL1_cfdna_ready | 164 bp | 0.3 X |
| 21564-PL1_cfdna_ready | 146 bp | 0.3 X |
| 21566-PL1_cfdna_ready | 139 bp | 0.4 X |
| 21569-PL0_cfdna_ready | 162 bp | 0.4 X |
| 21572-PL0_cfdna_ready | 159 bp | 0.5 X |
| 21580-PL1_cfdna_ready | 141 bp | 0.4 X |
| 21611-PL0_cfdna_ready | 167 bp | 0.2 X |
| 21614-PL0_cfdna_ready | 165 bp | 0.3 X |
| 21624-PL0_cfdna_ready | 168 bp | 0.4 X |
| 21627-PL0_cfdna_ready | 168 bp | 0.4 X |

#### Picard

Picard is a set of Java command line tools for manipulating high-throughput sequencing data.

##### Alignment Summary

Please note that Picard's read counts are divided by two for paired-end data. Total bases (including unaligned) is not provided.

Percentages | Aligned Reads | Aligned Bases | Export Plot

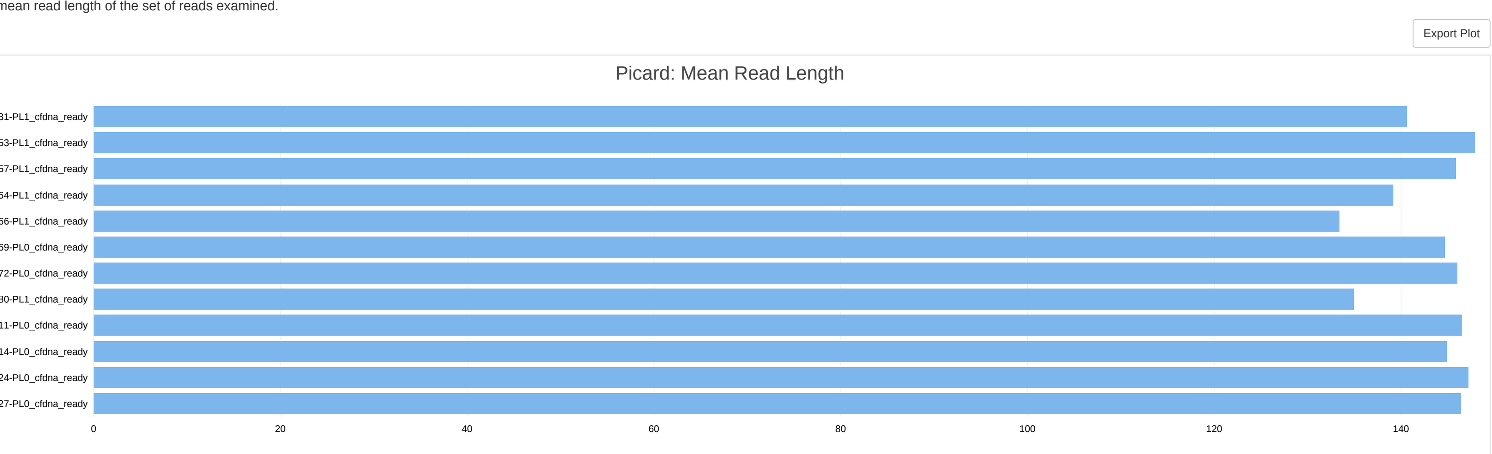


##### Mean read length

The mean read length of the set of reads examined.

Export Plot

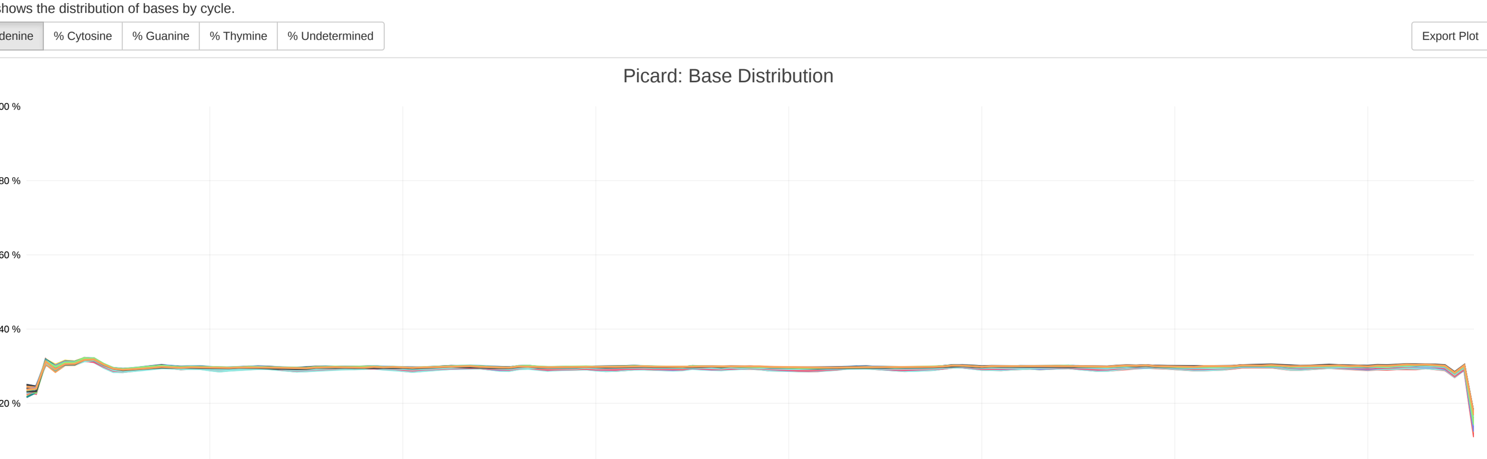


##### Base Distribution

Plot shows the distribution of bases by cycle.

% Adenine | % Cytosine | % Guanine | % Thymine | % Undetermined | Export Plot

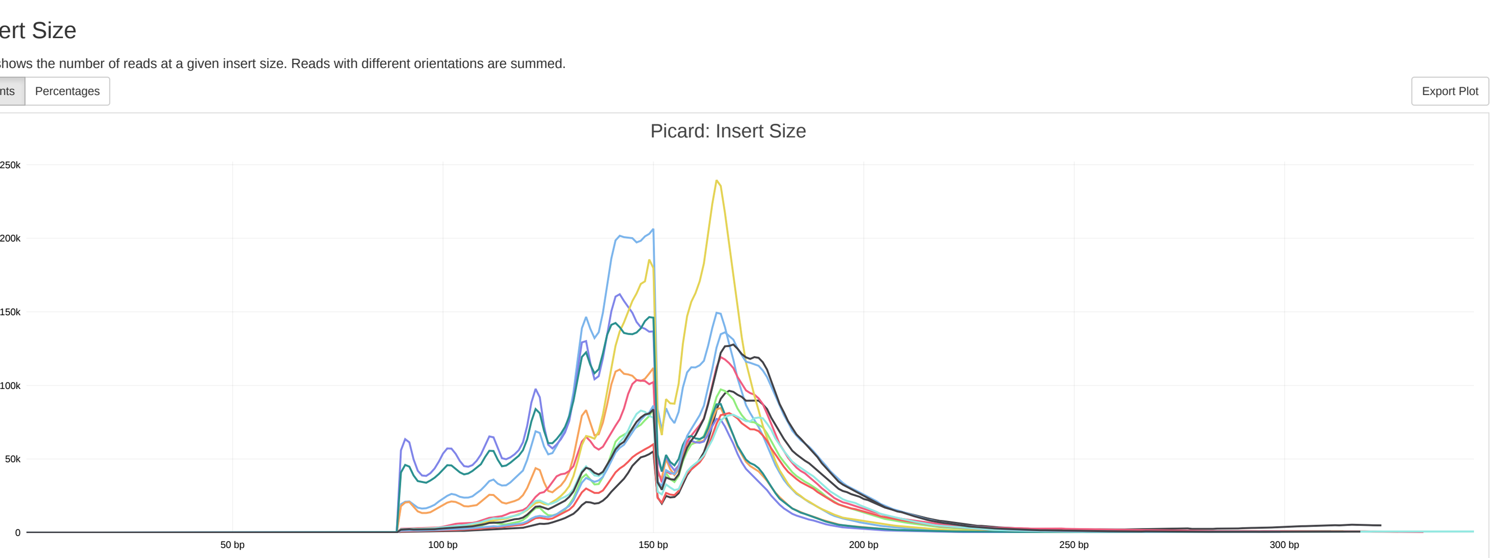


##### Insert Size

Plot shows the number of reads at a given insert size. Reads with different orientations are summed.

Counts | Percentages | Export Plot

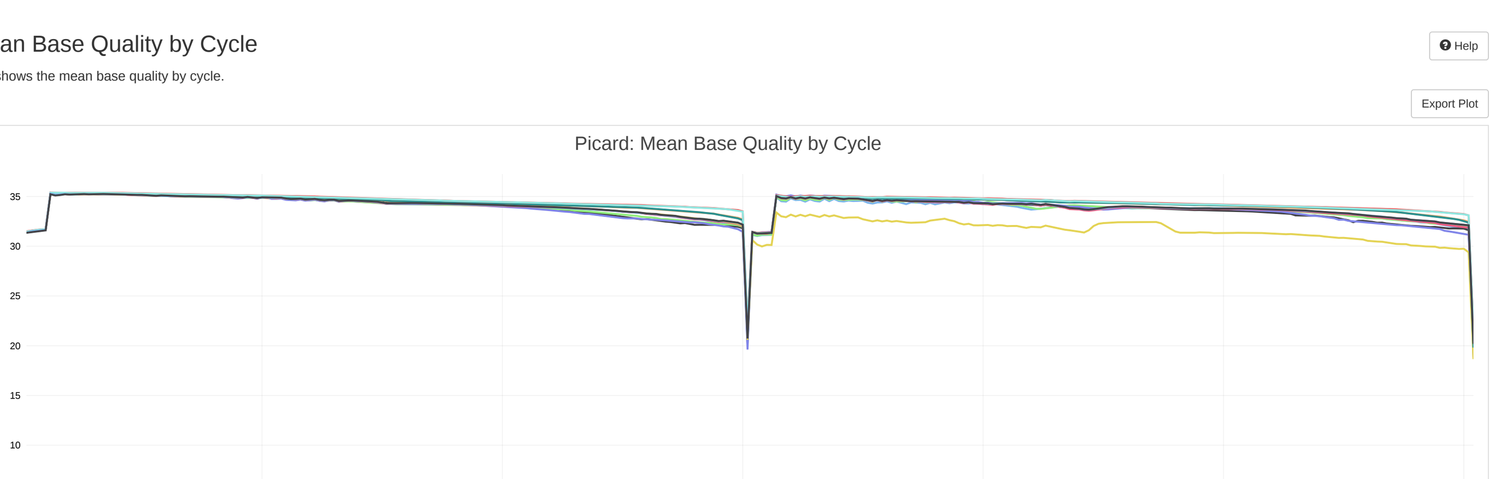


##### Mean Base Quality by Cycle

Help

Plot shows the mean base quality by cycle.

Export Plot

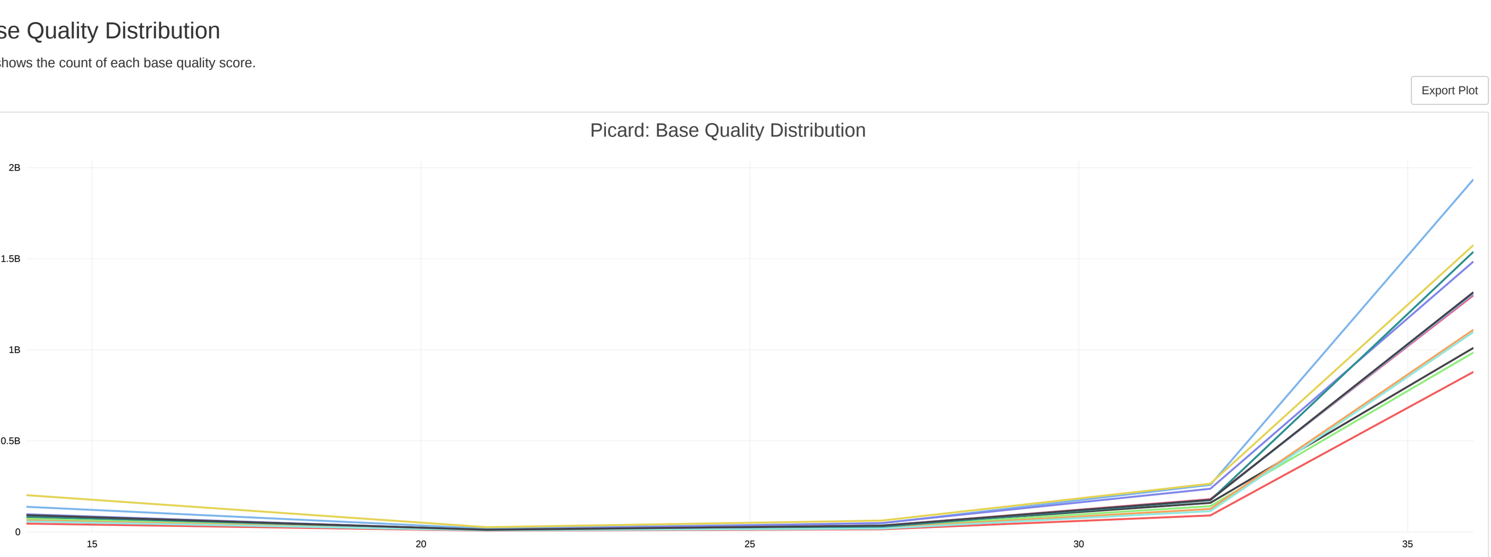


##### Base Quality Distribution

Plot shows the count of each base quality score.

Export Plot

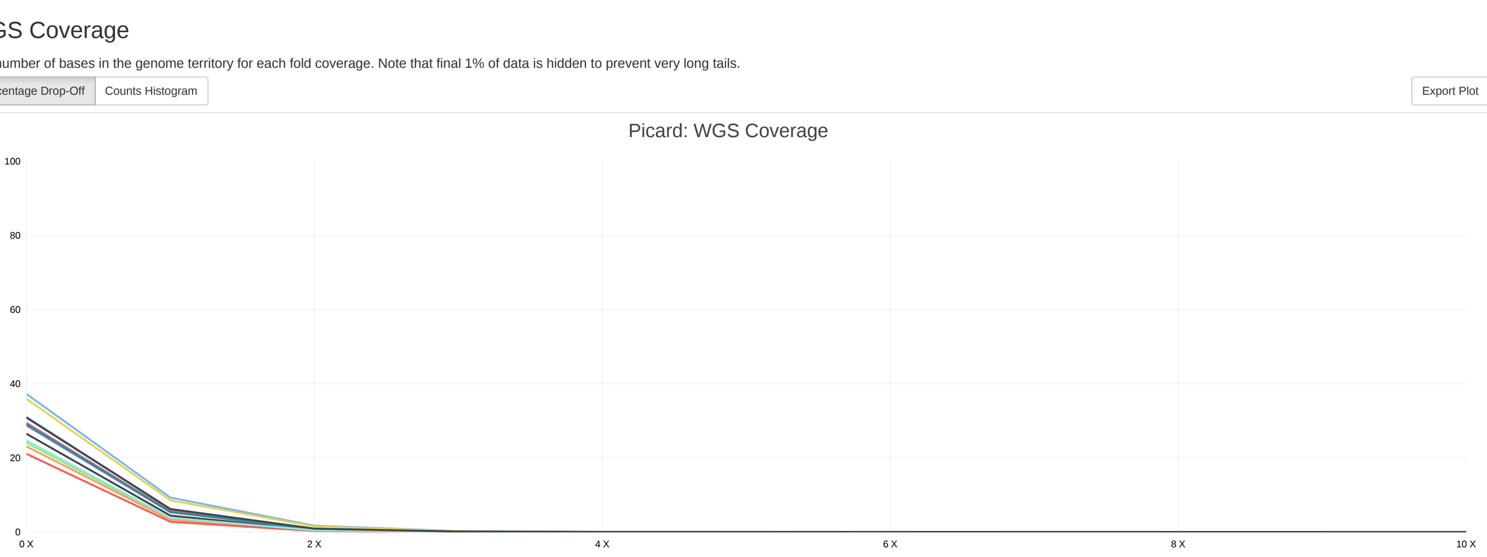


##### WGS Coverage

The number of bases in the genome territory for each fold coverage. Note that final 1% of data is hidden to prevent very long tails.

Percentage Drop-Off | Counts Histogram | Export Plot

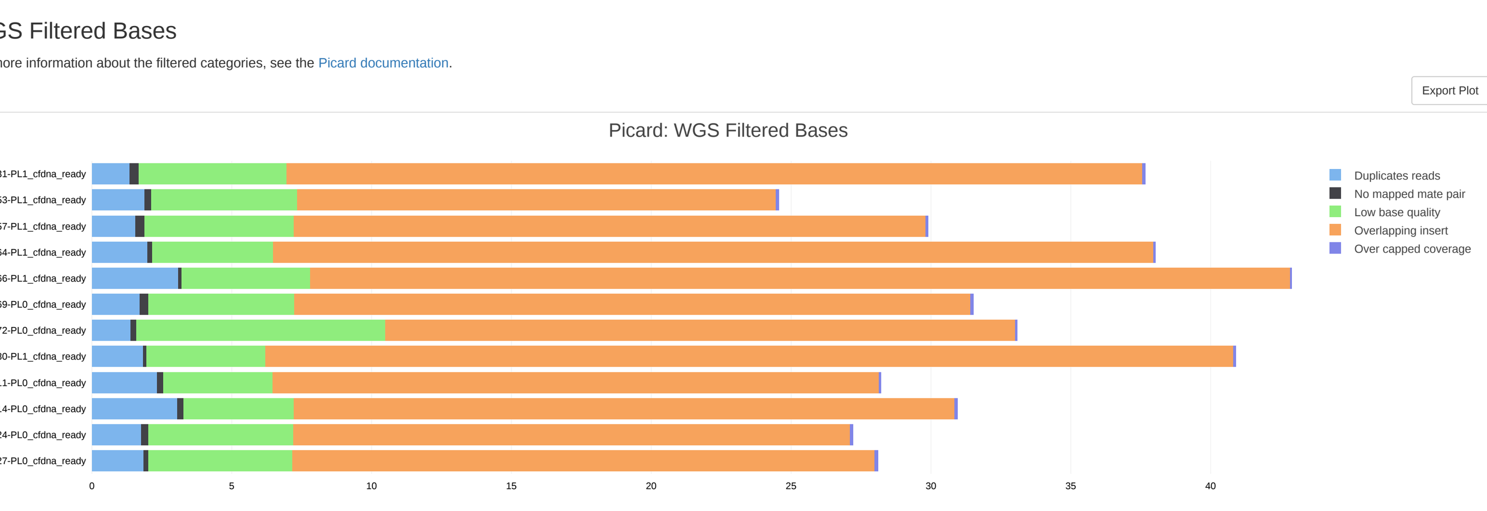


##### WGS Filtered Bases

For more information about the filtered categories, see the Picard documentation.

Export Plot

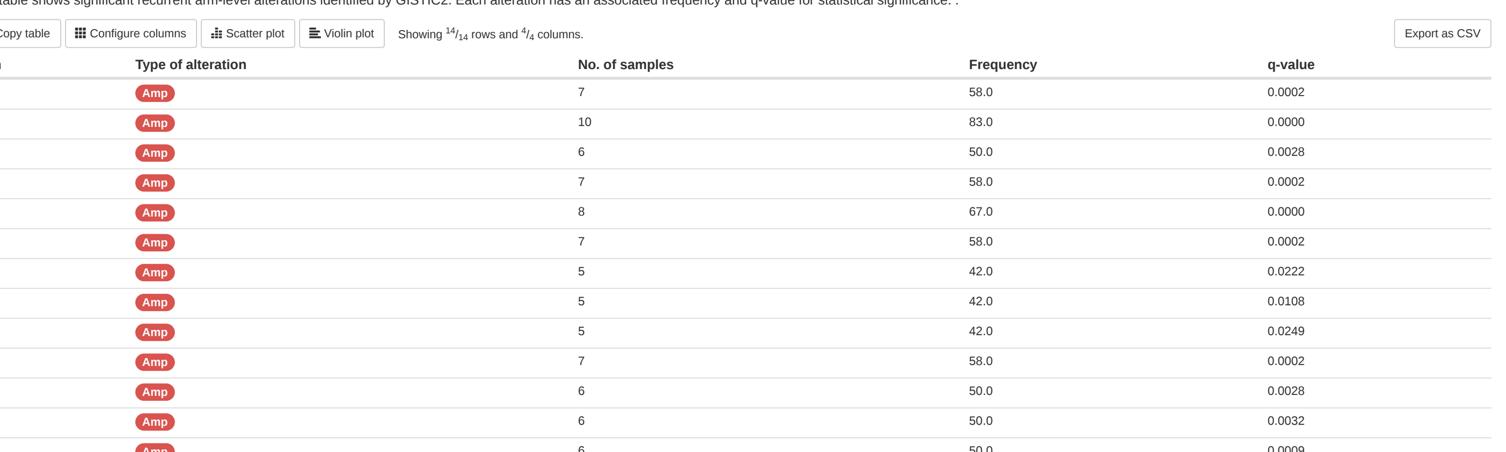


#### Gistic arm-level analysis summary

This table shows significant recurrent arm-level alterations identified by GISTIC2. Each alteration has an associated frequency and q-value for statistical significance. .

Copy table | Configure columns | Scatter plot | Violin plot | Showing 14/14 rows and 4/4 columns. | Export as CSV

| Arm | Type of alteration | No. of samples | Frequency | q-value |
|---|---|---|---|---|
| 1p | Amp | 7 | 58.0 | 0.0002 |
| 2p | Amp | 10 | 83.0 | 0.0000 |
| 3q | Amp | 6 | 50.0 | 0.0028 |
| 5p | Amp | 7 | 58.0 | 0.0002 |
| 6p | Amp | 8 | 67.0 | 0.0000 |
| 7q | Amp | 7 | 58.0 | 0.0002 |
| 8q | Amp | 5 | 42.0 | 0.0222 |
| 10p | Amp | 5 | 42.0 | 0.0108 |
| 11q | Amp | 5 | 42.0 | 0.0249 |
| 12p | Amp | 7 | 58.0 | 0.0002 |
| 19p | Amp | 6 | 50.0 | 0.0028 |
| 19q | Amp | 6 | 50.0 | 0.0032 |
| 20p | Amp | 6 | 50.0 | 0.0009 |
| 20q | Amp | 8 | 67.0 | 0.0000 |

#### IchorCNA

This table shows the tumor fraction and ploidy estimates made by ichorCNA.

Copy table | Configure columns | Scatter plot | Violin plot | Showing 12/12 rows and 3/3 columns. | Export as CSV

| Sample ID | Ploidy | Tumor fraction | MAD |
|---|---|---|---|
| 21614-PL1 | 2 | 0.04437 | 0.0530 |
| 21553-PL1 | 2 | 0.0419 | 0.0513 |
| 21557-PL1 | 2 | 0.08452 | 0.0510 |
| 21627-PL1 | 2 | 0.06006 | 0.0478 |
| 21611-PL1 | 2 | 0.1332 | 0.0584 |
| 21572-PL1 | 2 | 0.0922 | 0.0461 |
| 21624-PL1 | 2 | 0.1086 | 0.0488 |
| 21566-PL1 | 2 | 0.1358 | 0.0540 |
| 21569-PL1 | 2 | 0.1854 | 0.0528 |
| 21531-PL1 | 2 | 0.1291 | 0.0445 |
| 21580-PL1 | 2 | 0.1451 | 0.0531 |
| 21564-PL1 | 2 | 0.09609 | 0.0557 |

#### Gistic Analysis

Maftools genomic plot with segments highlighting significant Amplifications and Deletions.

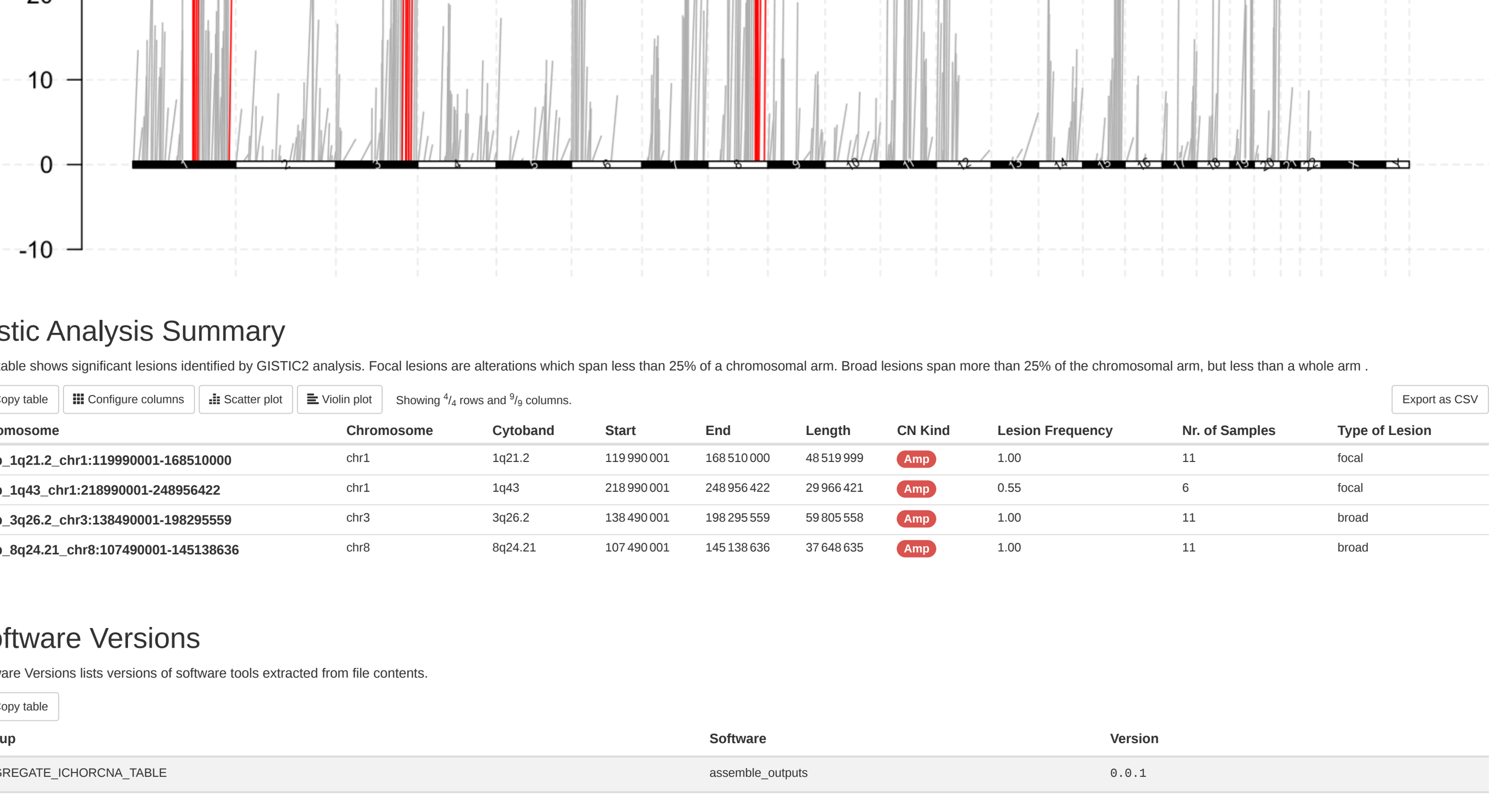


#### Gistic Analysis Summary

This table shows significant lesions identified by GISTIC2 analysis. Focal lesions are alterations which span less than 25% of a chromosomal arm. Broad lesions span more than 25% of the chromosomal arm, but less than a whole arm .

Copy table | Configure columns | Scatter plot | Violin plot | Showing 4/4 rows and 9/9 columns. | Export as CSV

| Chromosome | Chromosome | Cytoband | Start | End | Length | CN Kind | Lesion Frequency | Nr. of Samples | Type of Lesion |
|---|---|---|---|---|---|---|---|---|---|
| Amp_1q21.2_chr1:119990001-168510000 | chr1 | 1q21.2 | 119 990 001 | 168 510 000 | 48 519 999 | Amp | 1.00 | 11 | focal |
| Amp_1q43_chr1:218990001-248956422 | chr1 | 1q43 | 218 990 001 | 248 956 422 | 29 966 421 | Amp | 0.55 | 6 | focal |
| Amp_3q26.2_chr3:138490001-198295559 | chr3 | 3q26.2 | 138 490 001 | 198 295 559 | 59 805 558 | Amp | 1.00 | 11 | broad |
| Amp_8q24.21_chr8:107490001-145138636 | chr8 | 8q24.21 | 107 490 001 | 145 138 636 | 37 648 635 | Amp | 1.00 | 11 | broad |

#### Software Versions

Software Versions lists versions of software tools extracted from file contents.

Copy table

| Group | Software | Version |
|---|---|---|
| AGGREGATE_ICHORCNA_TABLE | assemble_outputs | 0.0.1 |
| ASSEMBLE_GISTIC_OUTPUT | gistic-cli | 0.4.2 |
| CONCATENATE_BIN_PLOTS | qpdf | 11.3.0 |
| CORRECT_LOGR_ICHORCNA | correct_logR_ichorcna | 0.1 |
| GISTIC2 | GISTIC2 | 0.1 |
| HMMCOPY_READCOUNTER_ICHORCNA | hmmcopy | 0.1.1 |
| MAFTOOLS | MAFTOOLS | 0.1 |
| PICARD_COLLECTMULTIPLEMETRICS | picard | 3.1.1 |
| RUN_ICHORCNA | ichorcna | 0.3.2 |
| Workflow | Nextflow | 24.4.2 |
| | dincalcilab/samurai | v1.0dev-gfd35d1f |

#### dincalcilab/samurai Methods Description

Suggested text and references to use when describing pipeline usage within the methods section of a publication.

##### Methods

Data was processed using dincalcilab/samurai v1.0dev of the nf-core collection of workflows (Ewels *et al.*, 2020).

The pipeline was executed with Nextflow v24.04.2 (Di Tommaso *et al.*, 2017) with the following command:

```
nextflow run DIncalciLab/swgscna -r master --input samplesheet.csv -params-file parameters.yaml --outdir output_ichorcna/ -profile cluster -qs 4
```


Notes:

- If available, make sure to update the text to include the Zenodo DOI of version of the pipeline used.
- The command above does not include parameters contained in any configs or profiles that may have been used. Ensure the config file is also uploaded with your publication!
- You should also cite all software used within this run. Check the "Software Versions" of this report to get version information.

#### dincalcilab/samurai Workflow Summary

- this information is collected when the pipeline is started.

**Core Nextflow options**

| | |
|---|---|
| revision | master |
| runName | pedantic_shirley |
| containerEngine | apptainer |
| launchDir | /home/incalci/lbeltrame/analysis/samurai-ichorcna |
| workDir | /home/incalci/lbeltrame/analysis/samurai-ichorcna/work |
| projectDir | /home/incalci/lbeltrame/.nextflow/assets/DIncalciLab/swgscna |
| userName | lbeltrame |
| profile | cluster |
| configFiles | N/A |

**Input/output options**

| | |
|---|---|
| input | samplesheet.csv |
| outdir | output_ichorcna/ |

**Reference genome options**

| | |
|---|---|
| fasta | /opt/bcbio/1.2.8/genomes/Hsapiens/hg38/seq/hg38.fa |
| fai | /opt/bcbio/1.2.8/genomes/Hsapiens/hg38/seq/hg38.fa.fai |
| dict | /opt/bcbio/1.2.8/genomes/Hsapiens/hg38/seq/hg38.dict |
| igenomes_ignore | true |

**Institutional config options**

| | |
|---|---|
| custom_config_base | /home/incalci/shared/pipeline-configs/ |

**Max job request options**

| | |
|---|---|
| max_cpus | 100 |
| max_memory | 260.GB |

**Generic options**

| | |
|---|---|
| hook_url | https://hookshot.heavensinferno.net/webhook/slack/a154b5f4-5a45-491c-9d4c-fffa71e7dd4d |
| multiqc_logo | /home/incalci/lbeltrame/.nextflow/assets/DIncalciLab/swgscna/assets/dincalcilab-samurai_logo_dark.png |

**Alignment options**

| | |
|---|---|
| aligner | false |

**Common sWGS options**

| | |
|---|---|
| caller | ichorcna |
| analysis_type | liquid_biopsy |

**Common liquid biopsy options**

| | |
|---|---|
| normal_panel | /home/incalci/shared/references/ctDNA_PoN_median.rds |

**ichorCNA options**

| | |
|---|---|
| ichorcna_readcounte… | chr1,chr2,chr3,chr4,chr5,chr6,chr7,chr8,chr9,chr10,chr11,chr12,chr13,chr14,chr15,chr16,chr17,chr18,chr19,chr20,chr21,chr22,chrX |
| ichorcna_chrs_to_use | paste0('chr', c(1:22, 'X')) |
| ichorcna_chrs_to_train | paste0('chr', c(1:22, 'X')) |
| ichorcna_chrs_to_no… | paste0('chr', c(1:22)) |
| ichorcna_max_cn | 6 |
| ichorcna_estimate_pl… | false |
| ichorcna_normal_stat… | 0.90, 0.98, 0.99, 0.995, 0.999 |
| ichorcna_gc_wig | /home/incalci/shared/references/ichorCNA/gc_hg38_500kb.wig |
| ichorcna_map_wig | /home/incalci/shared/references/ichorCNA/map_hg38_500kb.wig |
| ichorcna_centromere… | /home/incalci/shared/references/ichorCNA/GRCh38.GCA_000001405.2_centromere_acen.txt |

**GISTIC specific options**

| | |
|---|---|
| run_gistic | true |
| gistic_t_amp | 0.3 |
| gistic_t_del | 0.3 |
| gistic_remove_x | true |
| gistic_broad_analysis | true |
| gistic_broad_chr_len… | 0.7 |
